# The synovial lining macrophage layer develops in the first weeks of life in a CSF1- and TGFβ-dependent but monocyte-independent process

**DOI:** 10.1101/2025.05.21.655269

**Authors:** Marlene Magalhaes Pinto, Bert Malengier-Devlies, Guillaume Seuzaret, Anna Ahlback, Solvig Becker, Katelyn Patatsos, Georgios Drakoulis, Julia Karjalainen, Christiane Ruedl, David Voehringer, Calum C Bain, Elaine Emmerson, Barbora Schonfeldova, Kristina Zec, Irina Udalova, Theodoros Simakou, Lucy MacDonald, Mariola Kurowska-Stolarska, Jadwiga Zarebska, Tonia Vincent, Romeo Ricci, Eric Erbs, Jack Barrington, Barry W McColl, Georgiana Neag, Samuel Kemble, Christopher Mahony, Adam Croft, Louis Boon, Nicole Migotsky, Megan L Killian, Oumaima Ben Brahim, Stefan Uderhardt, Alexandre Gallerand, Stoyan Ivanov, Rebecca Gentek

## Abstract

Synovial joints harbor a protective lining layer, consisting of fibroblasts and macrophages, which form an epithelial-like barrier. In inflamed joints, lining macrophages regulate both early inflammatory cell influx and resolution. Despite these critical functions, it is currently unknown at what stage during development the synovial macrophage lining is established, and which signals drive this process. Here, we use a combination of genetic models and *in vivo* perturbations, single cell transcriptomics and imaging to delineate the process of lining formation in mice. We find that the synovial lining is immature at birth and becomes established within the first 3 weeks of life. In this window, the lining is gradually populated with macrophages that originate from fetal sources, proliferate and acquire the lining-specific transcriptional identity. In contrast, monocytes contribute only minimally to the developing lining, and their input remains limited in healthy adulthood. We identify CSF1 and TGFβ as key signals in this process, which also involves mechanosensing through PIEZO1. Our study thus identifies the early postnatal window as a critical period for lining macrophage development, with potential lifelong impact on joint health and disease.

## Introduction

Synovial joints contain a cavity filled with lubricating fluid, which enables friction-free movements between rigid skeletal elements. This cavity is lined with a specialised membrane that secretes the synovial fluid and forms a sterile barrier. Macrophages are the predominant immune cell present in the healthy synovium. Whilst their homeostatic functions have not fully been unravelled, synovial macrophages have been implicated in immune surveillance, debris clearing, recycling of synovial fluid components and synovial membrane integrity^1,2^. The synovial membrane comprises a lining layer and an adjacent sublining tissue, and distinct populations of macrophages reside in both compartments. The sublining is an adipose-like loose connective tissue, which is innervated and well vascularized and contains several different types of interstitial macrophages. The lining, on the other hand, is composed only of highly specialised fibroblasts and macrophages. Lining macrophages are transcriptionally distinct from other synovial macrophage populations, and they exhibit an anti-inflammatory phenotype and high levels of efferocytic receptors^1,3^. In mice, they are characterised by expression of VSIG4 (V-set and immunoglobulin domain containing 4) and the CX3CL1 (fractalkine) receptor CX3CR1^1^, which within joints is restricted to lining macrophages. Their human counterparts are TREM2^+^ MerTK^+^ macrophages^3^.

The synovial lining lacks a conventional basement membrane. Instead, it is held together by a dense extracellular matrix and lining macrophages, which intercalate with each other via epithelial-like cell-cell contacts^1^. These contacts are lost during acute joint inflammation, as lining macrophages rapidly change their morphology, orientation and phenotype^1^, allowing for inflammatory cell influx. At the early stages of joint inflammation, lining macrophages can also actively engulf antigen-immune complexes, which results in the release of chemo-attractants and neutrophil recruitment^4^. However, lining macrophages are also involved in resolution of inflammation at the later stages of (experimental) arthritis: they appear to retain their anti-inflammatory phenotype and remove dying neutrophils and monocytes^1^. Indeed, genetic depletion of lining macrophages prior to the onset of experimental arthritis exacerbates the severity of disease^1^, in line with an overall protective role. Lining macrophages therefore have an essential gatekeeping function and are central to joint health and disease.

Whilst recent studies have furthered our understanding of synovial lining macrophage biology, their development and ontogeny remain incompletely resolved. In adult joints, lining macrophages are thought to be largely monocyte-independent based on bone marrow chimeras, mice deficient in CCR2^5^, and short-term parabiosis^1^. However, it is unclear if this is also true under truly homeostatic conditions, and from which sources lining macrophages instead originate. Moreover, it is not known when and through which mechanisms the macrophage lining is first formed during development. These are important questions, because even after the resolution of synovial inflammation, many pre-existing macrophages appear to remain in the lining^1^. Furthermore, we increasingly recognise that the susceptibility to and severity of chronic inflammatory diseases including arthritis are enhanced by adverse environments in early life^6-10^. If lining macrophages are indeed long-lived and established in early life, then adverse conditions experienced during development may shape their functions long-term.

In this study, we comprehensively profiled the development of synovial macrophages, with a focus on the lining. We found that the lining is immature and only sparsely populated with macrophages in newborn mice and human foetuses. Through a combination of genetic fate mapping and transcriptomic profiling, we showed that in mice, the synovial macrophage lining is formed in the first 3 weeks of life from foetal-restricted progenitors. This process involves proliferation but is monocyte-independent, and monocyte contribution remains low during adult homeostasis. Development of the macrophage lining critically requires colony stimulating factor 1 (CSF1) and TGFβ signalling and may also involve sensing of mechanical loading through PIEZO1. Our study thus highlights the early postnatal period as the critical window for lining macrophage development.

## Results

### The synovial lining macrophage layer is formed in the first weeks of life

In mice, CX3CR1⁺ macrophages have been reported in the foetal joint from embryonic day (E) 15.5 onwards^1^. However, although cavitation has been initiated at this stage, there is no clear macrophage lining yet, and it was unknown when and how the macrophage lining develops. To gain first insights into how and when lining macrophages develop, we imaged synovial tissue from mouse knee joints from birth to adulthood, focusing on the meniscus region. In adult joints, lining macrophages form an epithelial-like barrier facing the joint cavity (Figure 1A). As previously described^1^, they express VSIG4 and CX3CR1, markers that within the joint are specific to this macrophage population. In stark contrast, although the cavity is already demarcated by fibroblasts (Figure S1A), the newborn joint lacks a continuous layer of lining macrophages (Figure 1A). At birth, macrophages can be found scattered throughout the joint, and those already present in the lining region are separated by gaps (Figure 1A, 1C) and lack VSIG4 expression (Figure 1D). The density of macrophages in the lining gradually increases over the first three weeks of life (Figure 1B) and the gaps are closed (Figure 1C), resulting in a continuous macrophage layer. In this period, lining macrophages also acquire VSIG4 (Figure 1D and 1E). CX3CR1, on the other hand, is expressed by all joint macrophages at birth and becomes progressively restricted to the lining population (Figure 1A). Similar to neonatal murine joints, we found that the synovial lining is immature in the human foetal knee (Figure S1B), indicating that this phenomenon is not species-specific. Macrophages are present in the lining region, but they do not form a continuous layer facing the cavity. However, we did observe an ∼1.7-fold increase in the macrophage density in the lining area between 13-16 and 17-19 weeks of gestation (Figure S1C), which might indicate the beginning of lining maturation. In addition to the maturation of the lining, we also observed changes in the sublining of the developing mouse joint. In adults, the sublining is a well-vascularized adipose tissue that also harbours several types of macrophages and fibroblasts^1^. Between birth and 3 weeks of age, adipocytes become lipid-filled and increase in size (Figure S1D, S1E). Joints are anatomically highly complex structures. Since our confocal imaging analyses focused on the meniscus region, we cannot exclude that these were localized effects. We thus applied high-resolution lightsheet microscopy on newborn joints to visualise lining macrophages in 3D (Figure 1F). Whilst CD44^+^ fibroblasts already cover the entire joint, macrophages do not form a continuous layer at the newborn stage. They are scattered throughout the joint instead but appear to localise in non-uniformly distributed clusters. These results confirm what we have observed in knee joint sections.

**Figure 1.**
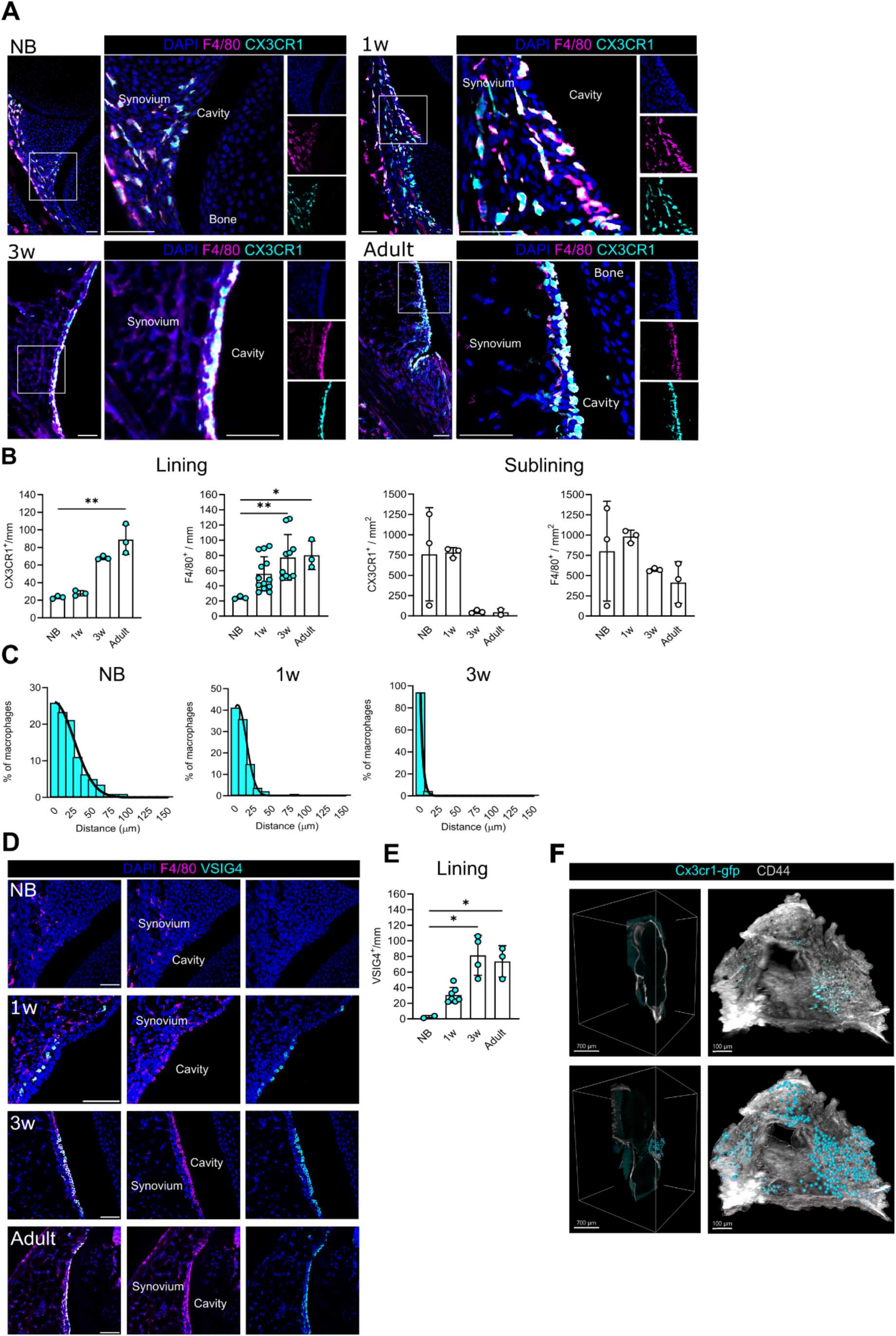
The synovial lining macrophage layer is formed in the first weeks of life. **(A)** Representative immunofluorescence images of synovial tissue from newborn (NB), 1-week-old (1w), 3-week-old (3w) and adult mice, stained for DAPI (blue), F4/80 (magenta), and further showing CX3CR1-GFP (cyan). Scale bars: 50 µm. **(B)** Quantification of total (F4/80^+^) or CX3CR1^+^ macrophage density in the synovial lining and sublining at the indicated developmental stages. Each dot represents an individual mouse, with quantification based on 2 to 3 sections per mouse. **(C)** Quantification of the distance between macrophages in the synovial lining area at the indicated ages. **(D)** Representative immunofluorescence images of synovial tissue stained for DAPI (blue), F4/80 (magenta), and VSIG4 (cyan). Scale bars: 50 µm. **(E)** Quantification of VSIG4⁺ macrophage density in the synovial lining across developmental stages. **(F)** High-resolution 3D lightsheet imaging of newborn murine knee joint of *Cx3cr1*^gfp^ mice stained for CD44 (grey) and showing Cx3cr1-GFP in cyan. Virtual sections and projected views are shown (Top). Automated spot detection highlights Cx3cr1-GFP⁺ macrophages (Bottom). Quantified data are presented as mean ± SD. Statistical significance was determined by Kruskal–Wallis test with Dunn’s correction for multiple comparisons, relative to newborn samples. *p<0.05, **p<0.01.

Collectively, these findings demonstrate that in mice, the macrophage lining is formed in the first weeks of life.

### Synovial macrophages are dynamic and lining macrophage identity is specified during postnatal development

To explore the cellular dynamics in the developing synovium in more depth and an unbiased manner, we next sought to perform single cell (sc)RNA-sequencing. First, we established that synovial macrophages can be isolated by micro-dissection and enzymatic digest of knee joint tissue^11^ from newborn to adult mice (Figure S2A). Flow cytometry showed a gradual increase in the proportion of joint macrophages expressing VSIG4 and a concomitant decrease in the relative abundance of CX3CR1^+^ macrophages (Figure 2A, 2B), consistent with our imaging data. We then isolated non-granulocytic immune cells (CD45^+^ Ly6G^-^) from knee joints of newborn, 1 week-old, 3-week-old and adult mice and subjected them to scRNA-sequencing on the 10x Chromium platform (Supplementary Table 3). In addition to immune cells, we also sequenced structural synovial cells, including fibroblasts (enriched as CD45^-^ PDPN^+^) and endothelial cells (CD45^-^ CD31^+^) to investigate tissue maturation processes and infer potential interactions between macrophages and structural cells of the synovium. Following quality filtering, integration and dimensionality reduction, unsupervised clustering analysis identified 19 distinct cell types, which we assigned manually using key transcriptional markers (Figure S2B, S2C, S2D, S2E). In addition to fibroblasts and endothelial cells, our synovial tissue samples also contained small populations of smooth muscle cells and muscle progenitors. Within the immune cell compartment, we found several types of macrophages as well as monocytes, dendritic cells, lymphocytes, mast cells and neutrophils (precursors). Neutrophils, neutrophil precursors, mast cells, and monocytes were most abundant in our newborn sample (Figure S2F), likely reflecting that although micro-dissected, early samples are more prone to contamination by surrounding tissues and blood.

**Figure 2.**
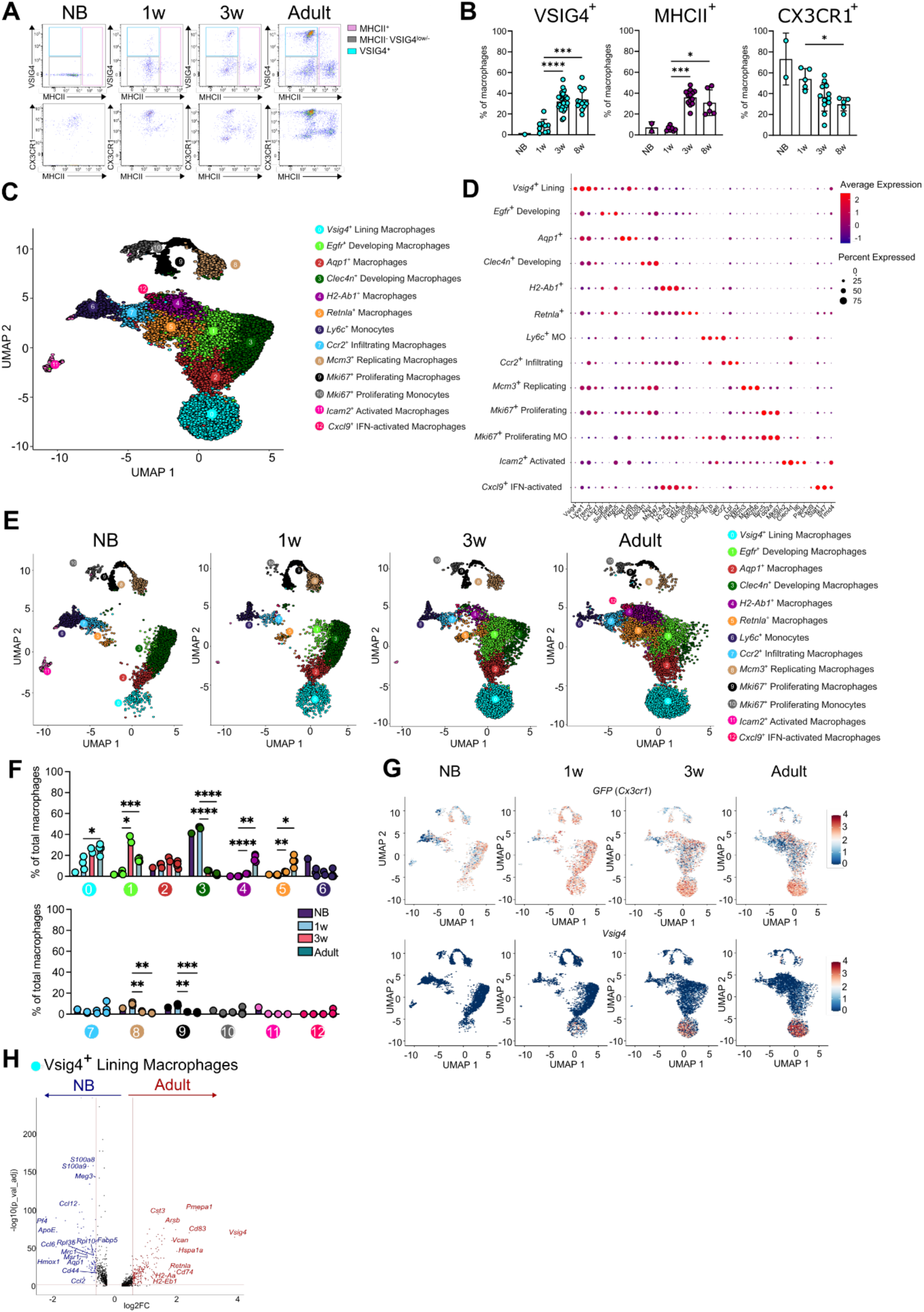
Lining macrophage identity is specified during postnatal development. **(A)** Representative flow cytometry plots of total macrophages (live CD45⁺ CD11b⁺ CD64⁺ F4/80⁺) in newborn (NB), 1-week-old (1w), 3-weeks-old (3w) and adult synovium, showing VSIG4 versus MHCII (top) and CX3CR1-GFP versus MHCII expression (bottom). **(B)** Flow cytometry quantification of VSIG4⁺, CX3CR1⁺, and MHCII⁺ macrophages at the indicated developmental stages. **(C)** Uniform manifold approximation and projection (UMAP) visualisation of single-cell RNA sequencing (scRNAseq) of synovial macrophages and monocytes across postnatal development. **(D)** Bubble plot showing pseudo bulk expression of key marker genes defining the different macrophage and monocyte clusters. **(E)** Time course UMAP visualisation of macrophage and monocyte populations at the indicated stages. **(F)** Quantification of macrophage and monocyte cluster distribution throughout development. **(G)** UMAP of lining macrophages overlaid with *(Cx3cr1-)gfp* and *Vsig4* expression. **(H)** Differentially expressed genes (DEGs) in newborn versus adult *Vsig4⁺* lining macrophages. For flow cytometry, each dot represents an individual mouse. For scRNAseq, each dot represents a mixed biological replicate as described in the Material and Methods section. Data are presented as mean ± SD. Statistical significance was determined using the Kruskal–Wallis test with Dunn’s correction for multiple comparisons, relative to 1w. *p<0.05, **p<0.01, ***p<0.001, ****p<0.0001.

To delineate macrophage dynamics during synovial tissue maturation at high resolution, we then proceeded to sub-clustering macrophages (*Apoe*, *Mrc1*, *Csf1r*) and monocytes (*Ly6c2*, *Il1b*, *Ccr2*). This yielded 2 monocyte and 11 macrophage clusters (Figure 2C, Figure S2G). Both monocyte clusters express *Ly6c2*, *Il1b*, and *Sell*. Proliferating monocytes (cluster 10) are characterized by expression of *Birc5*, *Top2a* and *Mki67*, whereas the other monocyte population (cluster 6) highly expresses *Clec4d*. Reassuringly, we found all macrophage populations that have previously been described in the ankle joint of adult mice^1^: *Vsig4*^+^ lining macrophages (cluster 0) (*Vsig4*, *Lyve1*, *Trem2*, *Cx3cr1*), *Aqp1*^+^ macrophages (cluster 2) (*Aqp1*, *Cd9*, *Cd109*), *MHCII*^+^ macrophages (cluster 4) (*H2-Aa*, *H2-Eb1*, *Cd74*), *Retnla*^+^ macrophages (cluster 5) (*Retnla*, *Ccl8*, *Cd209d*), and *Mki67*^+^ M-phase macrophages (cluster 9) (*Birc5*, *Top2a*, *Mki67*). Lastly, expression of *Ccr2*, *Lpl*, and elevated levels of *Il1b* distinguish a population of macrophages which is thought to be monocyte-derived and enriched in arthritic joints^1^ (cluster 7). In addition, we also found several synovial macrophage populations that have not previously been described. These are respectively defined by *Egfr* (cluster 1), *Clec4n* (cluster 3), markers associated with replication in S-phase (*Mcm3*, *Mcm4*, *Mcm6*) (cluster 8), expression of *Icam2* and a seemingly activated phenotype (*Clecd4*, *Il6*, *Padi4*) (cluster 11), and an IFN-signature and *Cxcl9* (*Cxcl9*, *Stat1*, *Ifi47*) (cluster 12). Of note, both *Icam2*^+^ and *Cxcl9*^+^ macrophages express high levels of *Timd4* (Figure 2D). We then analysed the prevalence of the different macrophage subtypes from neonates to adult mice. This uncovered distinct developmental patterns, which are comparable between mice of both sexes (Figure S2H). Several of the macrophage subsets found in the adult synovium only emerge during postnatal development (Figure 2E, 2F). Lining macrophages are initially sparse in the newborn joint and gradually increase during the first 3 weeks of life, after which their proportion remains stable. This is in keeping with our imaging and flow cytometry data. Similarly, *Egfr*^+^ macrophages are very rare until 3 weeks, but unlike those in the lining, they peak at 3 weeks. *Retnla*^+^ macrophages are sparse and *MHCII*^+^ macrophages are absent until 1 week of age, and both expand from 3 weeks onwards. Finally, *Cxcl9^+^* IFN-activated macrophages are only present in adult joints and remain very scarce. In contrast, other subtypes are restricted to or predominantly found at earlier developmental stages. *Clec4n*^+^ macrophages are the most abundant population in newborns, gradually decline after 1 week of age, and are rare in adult joints. *Icam2*^+^ activated macrophages are limited to the newborn stage and missing from the adult synovium. Lastly, other populations are present at all stages analysed, but some of these vary in abundance. Proliferating S-Phase *Mcm3*^+^ macrophages and *Mki67*^+^ M-phase macrophages, as well as monocytes, are enriched in newborn and 1-week-old joints. Likewise, *Ly6c*^+^ monocytes and *Ccr2*^+^ macrophages are present across development but are relatively more abundant in neonates. Intriguingly, *Aqp1*^+^ macrophages are found in similar frequencies across development.

Next, we assessed gene expression changes over time, focusing on lining macrophages. In agreement with our imaging and flow cytometry data, *Cx3cr1(gfp)* is initially expressed across synovial macrophage populations, but specifically retained by lining macrophages in adults (Figure 2G). *Vsig4*, on the other hand, is exclusive to lining macrophages at all stages, but only gradually acquired during postnatal maturation (Figure 2G). To better understand their maturation process, we then conducted differential gene expression analysis comparing adult lining macrophages with those already present at birth (Figure 2H). This revealed substantial transcriptional changes, with 271 genes up- and 160 genes downregulated. In addition to *Vsig4*, expression of *Trem2* is upregulated during maturation, another cardinal marker of mature lining macrophages. Increased expression was also observed in MHCII-related genes (*Cd74*, *H2-Ab1*, *H2-Aa*, *Cd83*, *Cst3*), genes involved in SMAD binding and TGFβ signalling (*Elmo1*, *Pmepa1*, *Skil*, *Mef2a*, *TgSr1*), cell adhesion (*Itgav*, *Fgfr1*, *F11r*), actin binding (*Myo1e*, *Sptbn1*, *Cald1*, *Myh9*) and GTPase activity (*Arhgap22*, *Rgs10*, *Gnaq*, *Rasgef1b*). Intriguingly, *Aqp1* is downregulated in the process. Other downregulated genes are associated with ribosomal function (*Rps27a*, *Rpl29*, *Rpl32*, *Rpl15*), chemokines (*Ccl12*, *Pf4*, *Ccl6*, *Ccl9*, *Ccl2*) and mannose or carbohydrate binding (*Clec4n*, *Clec4a2*, *Mrc1*). Transcriptomic differences remain pronounced for lining macrophages at 1 week and 3 weeks compared to adult joints (Figure S2I), but the number of DEGs gradually decreases, indicating an ongoing maturation process. Of note, apart from sex-linked genes, lining macrophages are transcriptionally comparable between male and female mice at 1 week, 3 weeks and in adulthood (Figure S2J).

In conclusion, we uncovered dynamic changes in the composition of the synovial macrophage compartment during postnatal development and demonstrated that lining macrophages are specified during the first weeks of life.

### The synovial lining is populated by foetal-derived macrophages with limited monocyte contribution

Bone marrow chimeras, short-term parabiosis (6 weeks) and CCR2 reporter mice indicate limited turnover of adult synovial lining macrophages from bone marrow monocytes^1,12,13^. By exclusion, it has therefore been suggested that they originate from foetal-restricted progenitors and self-maintain^1^. However, this has not formally been demonstrated, and the contribution of monocytes to the developing lining has also not been investigated. The precise developmental origin and kinetics of synovial lining macrophages thus remains unclear. To delineate their ontogeny, we thus conducted fate mapping experiments with a range of complementary models. First, we determined to what extent granulocyte-monocyte progenitor (GMP)-derived monocytes give rise to lining macrophages using *Ms4a3*^Cre^:*Rosa26*^lsl-tdT^:*Cx3cr1*^gfp^ mice, which allow simultaneous detection of CX3CR1 expressing cells (GFP) and cells deriving from GMP-dependent monocytes (tdTomato; TdT) ^14^. Confocal imaging showed that monocyte contribution to lining macrophages does not exceed on average 10% between birth and 3 weeks (Figure 3A), although the lining is populated with macrophages in this window. We then performed an extended longitudinal analysis up to 1 year using flow cytometry (Figure 3B). This revealed that the lining incorporates a small pool of monocytes between 3 and 6 weeks, remains stable until 6 months, and increases to ∼25% in 1-year-old mice. MHCII^+^ and MHCII^-^ sublining macrophages, on the other hand, showed more labelling at all stages, demonstrating monocytic origin. Because we intended to study the origins of synovial macrophages, our scRNAsequencing analysis was carried out on *Ms4a3*^Cre^:Rosa26^lsl-tdT^:*Cx3cr1*^gfp^ mice. The distribution of the *tdTomato* transcript confirmed our imaging and flow cytometry-based findings. As expected, almost all monocytes (Figure 3C, 3D) as well as neutrophils and neutrophil precursors (Figure S3A, S3B) express *tdTomato* transcript. Similarly, *Icam2*^+^ and *Ccr2*^+^ macrophages showed near-complete coverage with *tdTomato* signal (Figure 3C, 3D), in keeping with the notion that they are infiltrating populations^1^. From newborns to adults the frequency of *tdTomato*^+^ cells increased for *MHCII*^+^ and *Retnla*^+^ macrophages but remained low for *Vsig4*^+^ lining macrophages (∼14% in adult mice). Intriguingly, we noted that adult males had more monocyte-derived *tdTomato*^+^ macrophages in all clusters compared to females (Figure S3C), a finding that we also confirmed in an independent cohort of mice (Figure S3D). We then compared gene expression profiles of *tdTomato*^+^ and *tdTomato*^-^ macrophages within key synovial macrophage populations (Figure S3E). Overall, monocyte-derived and monocyte-independent macrophages are transcriptionally very similar, in line with the consensus that macrophage phenotype and function are predominantly imprinted by the microenvironment or niche, rather than their developmental origin^15–17^. However, we did observe subtle transcriptomic differences. Within *Vsig4*^+^ lining as well as *Aqp1*^+^, *Clec4n*^+^ and *Egfr*^+^ macrophages, the *tdTomato*^+^ fraction appears to have a more activated phenotype, with higher expression of MHCII-related genes (*H2-Ab1*, *H2-Eb1*, *H2-Aa*, and *Cd74*) and *Ccl8*, and lower levels of the angiogenic inducer *Ccn1*. For the *MHCII*^+^ and *Retnla*^+^ populations, *tdTomato*^+^ macrophages show increased *Ccr2* expression and lower levels of *Cd209* and *Timd4*, consistent with the growing consensus that Tim4 expression is increased with tissue residency.

**Figure 3.**
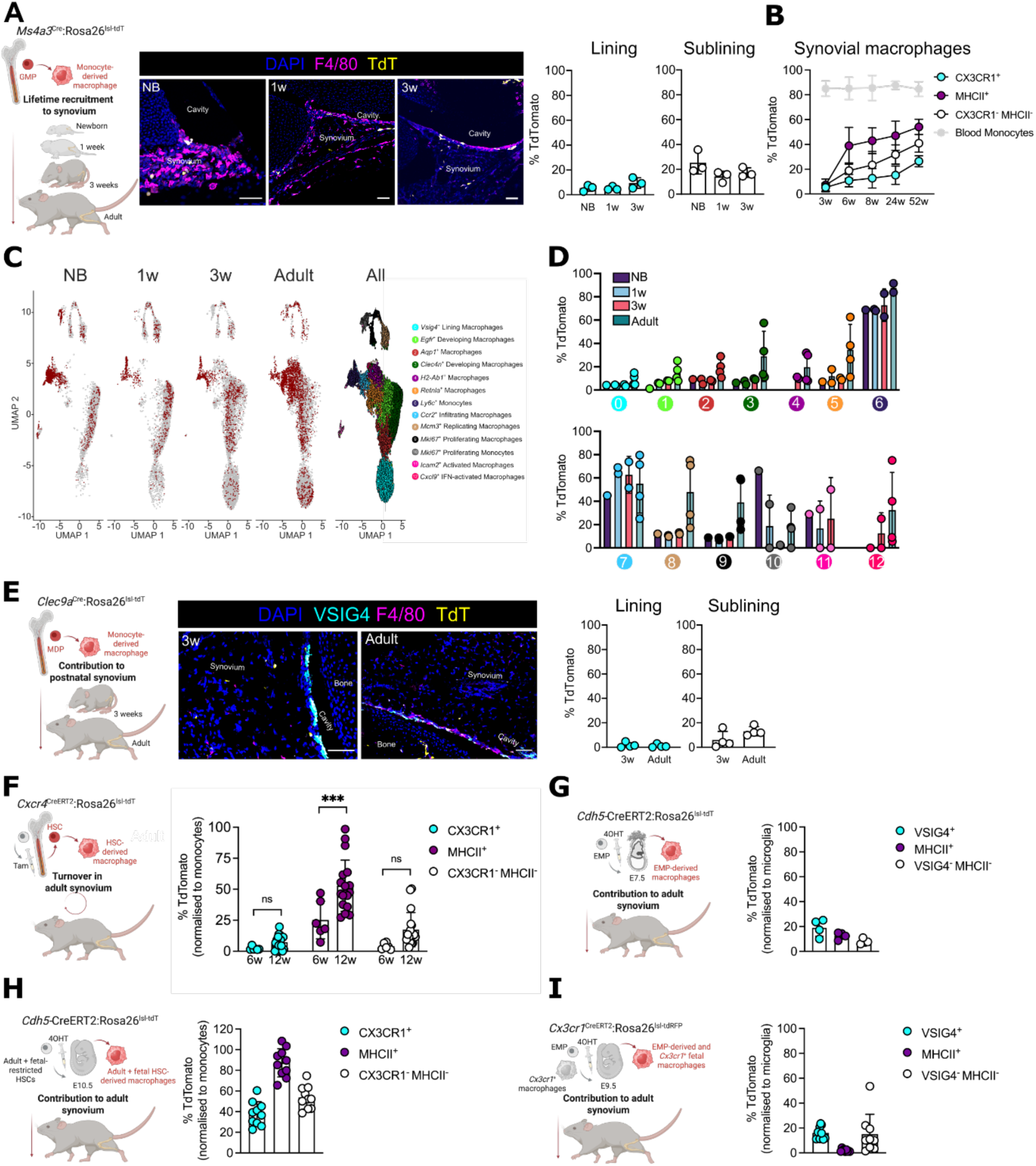
The synovial lining is populated by foetal-derived macrophages with limited monocyte contribution. (A-D) Contribution of granulocyte-monocyte progenitor (GMP)-derived monocytes to synovial macrophages. **(A)** (Left) Schematic of the *Ms4a3*^Cre^:Rosa26^lsl-tdT^ fate mapping model, which labels granulocyte-monocyte progenitor (GMP)-derived cells. (Middle) Representative immunofluorescence images of synovial tissue from *Ms4a3*^Cre^:Rosa26^lsl-tdT^ mice at the newborn (NB), 1-week (1w), and 3-weeks-old (3w) stages, showing tdTomato (tdT, yellow) and staining for DAPI (blue), F4/80 (magenta). (Right) Quantification of tdTomato⁺ macrophages in the synovial lining and sublining across developmental stages. **(B)** Time course analysis of monocyte contribution to synovial macrophages. Flow cytometry was used to quantify tdTomato⁺ cells within CX3CR1⁺ lining macrophages and CX3CR1⁻ MHCII⁻ and MHCII⁺ sublining macrophages from 3 weeks to 1 year of age. Ly6C^high^ blood monocytes served as a positive control. **(C)** Uniform manifold approximation and projection (UMAP) overlay of single-cell RNA sequencing (scRNAseq) showing *tdTomato* expression across all macrophage and monocyte clusters. **(D)** scRNAseq quantification of *tdTomato*⁺ macrophage subsets at the indicated life stages. **(E)** Contribution of monocyte and dendritic cell progenitors (MDPs) to synovial macrophages. (Left) Scheme depicting the *Clec9a*^Cre^:Rosa26^lsl-tdT^ fate mapping model, which labels cells originating from MDPs. (Middle) Representative immunofluorescence images of synovial tissue from *Clec9a*^Cre^:Rosa26^lsl-tdT^ mice at 3 weeks and in adulthood, showing and tdTomato (yellow) and stained for DAPI (blue), VSIG4 (cyan), F4/80 (magenta). (Right) Quantification of tdTomato⁺ macrophages in the synovial lining and sublining at the indicated ages. **(F)** Turnover of synovial macrophages from bone marrow haematopoietic stem cells (HSCs). (Left) Illustration of the *Cxcr4-* CreERT2:Rosa26^lsl-tdT^ fate mapping model. In adult mice, tamoxifen administration labels HSCs and their progeny in a pulse-chase manner. (Right) Flow cytometry analysis of tdTomato labelling in CX3CR1⁺ lining and CX3CR1⁻ MHCII⁻ and CX3CR1⁻ MHCII⁺ sublining populations 6 and 12 weeks post injection. **(G)** Contribution of yolk sac erythro-myeloid progenitors (EMPs) to synovial macrophages. (Left) Schematic of the *Cdh5*^CreERT2^:Rosa26^lsl-tdT^ fate mapping model. A single dose of 4-hydroxy-tamoxifen (4OHT) administered at embryonic day (E)7.5 labels EMPs and their progeny. (Right) Quantification of tdTomato⁺ labelling within CX3CR1⁺ lining and CX3CR1⁻ MHCII⁻ and CX3CR1^-^ MHCII⁺ sublining macrophages. Flow cytometry was performed on adult joints. **(H)** Contribution of later progenitors to synovial macrophages. (Left) In *Cdh5*^CreERT2^:Rosa26^lsl-tdT^ mice, 4OHT induction at E10.5 labels adult-type and foetal-restricted HSCs and their progeny. (Right) Flow cytometric quantification of tdTomato⁺ labelling in CX3CR1⁺ lining and CX3CR1⁻ MHCII⁻ and CX3CR1^-^ MHCII⁺ sublining macrophages of adult joints. **(I)** Persistence of EMP-derived (pre-)macrophages in the adult joint. (Left) Illustration of the *Cx3cr1*^CreERT2^:Rosa26^lsl-tdRFP^ model, in which 4OHT treatment at E9.5 labels *Cx3cr1*-expressing (pre-)macrophages and myeloid progenitors. (Right) Flow cytometric quantification of tdRFP labelling in the adult VSIG4⁺ lining and VSIG4⁻ MHCII⁻ and VSIG4^-^ MHCII⁺ sublining compartments. Scale bars: 50 µm. For imaging analyses, each dot represents a single animal, with quantification based on 2 to 3 sections per mouse. For flow cytometry, each dot represents an individual mouse. For scRNAseq, each dot represents a mixed biological replicate as described in the Material and Methods section. Data are presented as mean ± SD. Statistical significance was determined using the Mann–Whitney test for pairwise comparisons. ***p<0.001.

Monocytes themselves can have distinct origins. Those originating from GMP and monocyte-dendritic cell progenitors (MDP) are functionally different^18^, and MDP-derived monocytes also produce specific progeny, such as CD226^+^ adipose tissue macrophages, which are seeded in early life^19^. It has been proposed that MDP-derived monocytes and their macrophage progeny are labelled in *Clec9*^Cre^:Rosa26^lsl-tdT^ mice^19^. However, we found no labelling of synovial lining macrophages at 3 weeks and in adult joints of *Clec9*^Cre^:Rosa26^lsl-tdT^ mice (Figure 3E), and only low labelling (on average 10%) in sublining macrophages. The synovial lining macrophage layer thus forms independently of MDP-derived monocytes.

*Ms4a3*^Cre^ and *Clec9a*^Cre^ both constitutively label precursors and therefore provide a readout for lifetime monocyte recruitment (i.e. until the age of analysis). However, they do not delineate the time of recruitment, nor do they allow one to quantitate homeostatic turnover once adulthood is reached. To address these aspects of lining macrophage ontogeny, we made use of a number of tamoxifen-inducible fate mapping models. Hematopoietic stem cells (HSCs) express *Cxcr4* and *Kit*. Administration of tamoxifen to adult *Cxcr4*^CreERT2^:Rosa26^lsl-tdT^ or *Kit*^MerCreMer^:Rosa26^lsl-eYFP^ mice irreversibly labels HSCs and their progeny, and these approaches can therefore be used to determine the degree of turnover from the bone marrow in a pulse chase manner^20,21^. In tamoxifen-treated *Cxcr4*^CreERT2^:Rosa26^lsl-tdT^ mice, circulating monocytes were nearly fully labelled as anticipated and hence used for normalization. Synovial lining macrophages exhibited maximally 25% labelling at 12 weeks, whereas more than half of the MHCII^+^ sublining macrophages were labelled within this window, and the CX3CR1^-^ MHCII^-^ population showed intermediate levels of labelling (Figure 3F). These findings were corroborated in the *Kit*^MerCreMer^ model, in which CX3CR1^+^ lining macrophages showed minimal labelling 4 months after the tamoxifen pulse, whereas the majority of MHCII^+^ sublining macrophages was labelled (Figure S3F). Together, data obtained in monocyte- and HSC-fate mapping models demonstrated that at steady state, synovial lining macrophages develop and maintain with minimal BM monocyte contribution.

To address if synovial lining macrophages instead originate from fetal-restricted progenitors, we first used *Cdh5*^CreERT2^:Rosa26^lsl-tdT^:*Cx3cr1*^gfp^ mice. In this mouse line, distinct hematopoietic waves can be labelled according to the time of induction by 4-Hydroxy-tamoxifen (4OHT)^22^. A single dose of 4OHT delivered at E7.5 labels yolk sac (YS) erythro-myeloid progenitors (EMPs) and fate maps their progeny, whereas induction at E10.5 labels a combination of adult-type HSCs that colonise the bone marrow and the less well-defined “foetal-restricted HSCs”^22–25^. In adult mice pulsed at E7.5, lining macrophages were labelled at approximately 20% (Figure 3G). Macrophages originating from early YS EMPs thus constitute a fraction of the lining population. The majority thus derives from a later hematopoietic source. Indeed, on average 40% of lining macrophages were labelled in adult *Cdh5*^CreERT2^:Rosa26^lsl-tdT^:*Cx3cr1*^gfp^ mice pulsed at E10.5 (Figure 3H). Of note, MHCII^-^ sublining macrophages showed a similar degree of labelling, and MHCII^+^ sublining macrophages were labelled at nearly 90%. Whilst this approach cannot formally delineate the relative contribution of adult-type and foetal HSCs, we have shown conclusively in the *Ms4a3*^Cre^, *Clec9*^Cre^, *Cxcr4*^CreERT2^ and *Kit*^MerCreMer^ models that the lining receives limited input from BM progenitors. Labelling of lining macrophages following E10.5 induction in the *Cdh5*^CreERT2^ model is thus most consistent with these lining macrophages originating from fetal-restricted HSCs. Lastly, we opted for *Cx3cr1*^CreERT2^ mice as a complementary approach. Here, label induction at E9.5 fate maps EMP-derived committed “pre-macrophages”^26^ and the first terminally differentiated foetal macrophages, both of which express CX3CR1 at this stage^27,28^. In adult joints of *Cx3cr1*^CreERT2^:Rosa26^lsl-tdRFP^ mice pulsed at E9, ∼15% of lining macrophages were labelled (Figure 3I), indicating that these originate from EMP-derived (pre-)macrophages. YS EMPs thus contribute to the first macrophages populating the synovial lining, but these become superseded by later hematopoietic sources that are distinct from adult-type HSCs producing monocytes.

In summary, complementary fate mapping approaches showed that synovial lining macrophages develop in the first weeks of life and originate from foetal-restricted progenitors with minimal contribution from BM monocytes. This resembles lung alveolar macrophages^29^ and epidermal Langerhans cells^30^, which are also both seeded during a restricted developmental window.

### Postnatal expansion of lining macrophages is facilitated by proliferation and may involve *Aqp1*^+^ intermediates

We next wanted to delineate the mechanisms by which synovial lining macrophages expand postnatally, as well as their relationships with other synovial macrophage subsets. To investigate whether the lining population expands from macrophages already present in the neonatal joint, we leveraged the finding that all synovial macrophages uniformly express CX3CR1 at birth (Figure 1). In adult *Cx3cr1*^CreERT2^:Rosa26^lsl-tdRFP^ mice that had received tamoxifen as newborns, most (∼ 87%) lining macrophage were fate mapped at 3 weeks and more than half of these remained labelled in adult joints. MHCII^-^ sublining macrophages were labelled to a slightly lower degree (∼ 72% at 3 weeks and 32% in adults), and MHCII^+^ were only labelled to 34% at 3 weeks and declined to 9% in adults (Figure 4A). Lining macrophages thus expand from cells that express CX3CR1 at birth, which includes differentiated macrophages.

**Figure 4.**
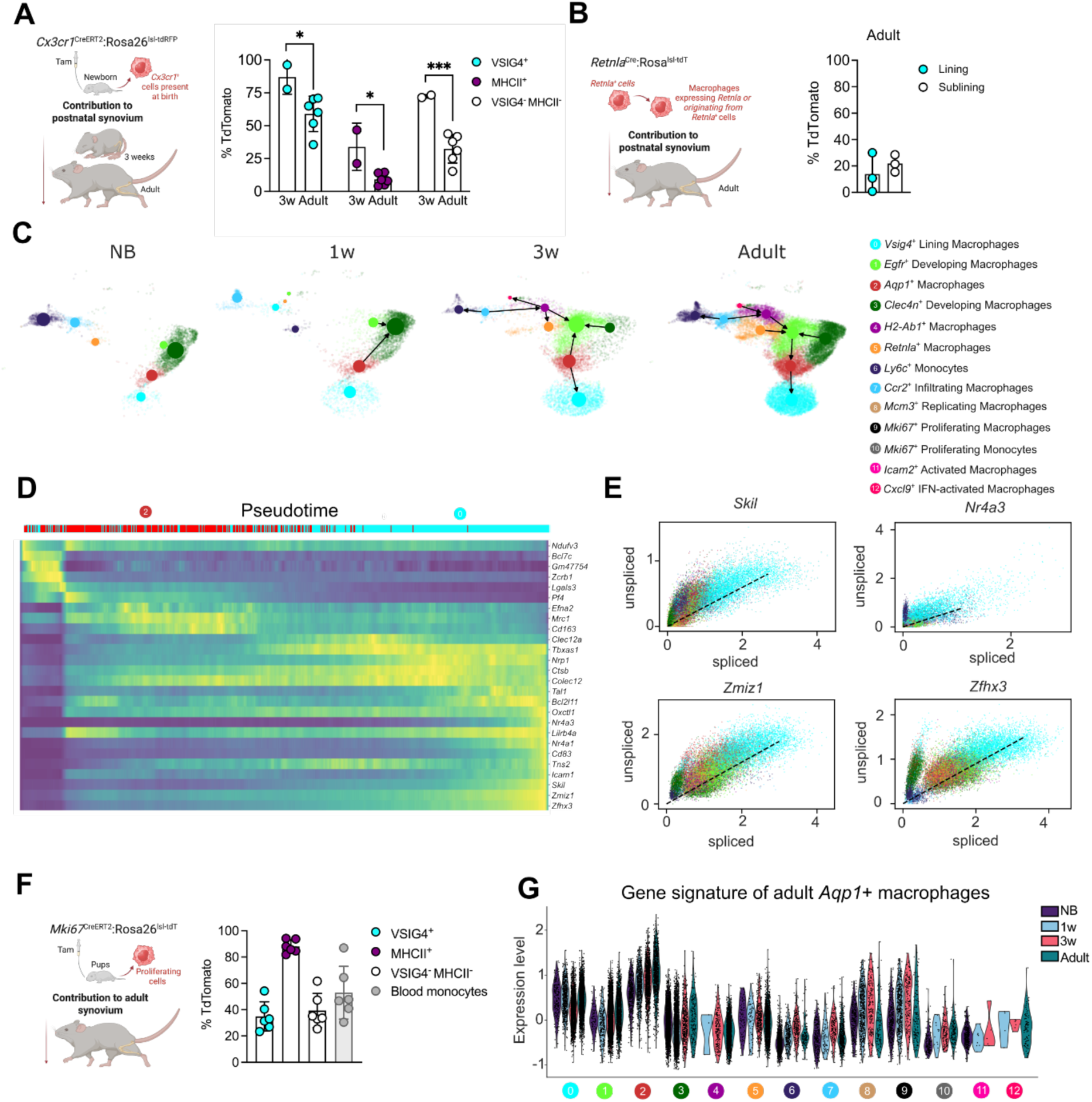
Lining macrophages expand postnatally through proliferation of pre-existing cells likely involving *Aqp1*^+^ intermediates. (A) Persistence of neonatal macrophages in synovial tissue. (Left) Schematic of the *Cx3cr1*^CreERT2^:Rosa26^lsl-tdRFP^ fate mapping model used to label *Cx3cr1*-expressing cells in neonates. (Right) Flow cytometric quantification of tdRFP labelling in VSIG4⁺ lining and VSIG4⁻ MHCII⁻ and VSIG4^-^ MHCII⁺ sublining macrophages at 3 weeks (3w) and in adulthood. **(B)** Contribution of *Retnla*^+^ cells to synovial macrophages. (Left) Scheme depicting the *Retnla*^Cre^:Rosa26^lsl-tdT^ model, which labels cells with current or a history of *Retnla* expression. (Right) Immunofluorescence-based quantification of tdTomato⁺ macrophages in the lining and sublining of adult joints. **(C)** Computational inference of developmental relationships between synovial macrophage populations. Partition-based graph abstraction (PAGA) plot overlaid onto uniform manifold approximation and projection (UMAP) embedding, showing RNA velocity trajectories among macrophage populations in newborn (NB), 1-week-old (1w), 3-weeks-old (3w) and adult synovium. **(D)** Transcriptional shift during the predicted differentiation of *Aqp1*⁺ intermediates to *Vsig4*⁺ lining macrophages. **(E)** Identification of potential factors driving lining macrophage differentiation based on RNA velocity pseudotime analysis. **(F)** Contribution of cells proliferating in the neonatal period to adult synovial macrophages. (Left) Schematic of the *Ki67*^CreERT2^:Rosa26^lsl-tdT^ fate mapping model. Tamoxifen was used to label proliferating cells in neonates. (Right) Flow cytometric quantification of tdTomato labelling in VSIG4⁺ lining and VSIG4⁻ MHCII⁻ and VSIG4^-^ MHCII⁺ sublining macrophages in adult synovium. **(G)** Expression of a gene set characteristic of adult *Aqp1*^+^ macrophages across synovial macrophage subsets and developmental stages. For flow cytometry, each dot represents an individual mouse. Data are presented as mean ± SD. Statistical significance was determined using the Mann– Whitney test for pairwise comparisons. *p<0.05, ***p<0.001.

It has previously been proposed that one source for lining macrophages in adult joints is *Csf1r*-expressing sublining macrophages^1^. A population of these sublining macrophages are defined by expression of *Retnla*. We therefore took advantage of *Retnla*^Cre^:Rosa26^lsl-tdT^ mice, in which active and historic expression of *Retnla* can be tracked through irreversible tdTomato labelling. This system showed that only ∼12% of synovial lining macrophages were labelled in adult *Retnla*^Cre^:Rosa26^lsl-tdT^ mice (Figure 4B), confirming that *Retnla*^+^ cells are not a major source for lining macrophages at any point in development. We then applied trajectory analyses to our single cell transcriptomic data to infer which of the other synovial macrophage populations might give rise to lining macrophages. RNA velocity analysis, supported by a tool which evaluates the confidence of cell transition states (PAGA), suggested there is minimal transition between populations in the newborn data (Figure 4C). However, in the 3-weeks and adult datasets, *Aqp1*^+^ macrophages were predicted as key branching points giving rise to *Vsig4*^+^ lining macrophages. A similar pathway to supplementing lining macrophages has recently been identified by Schonfeldova and colleagues in adult joints^31^. Our scRNAseq data suggest that the *Aqp1*^+^ population is also a potential local precursor to synovial lining macrophages during postnatal development. To elucidate possible drivers that might mediate the differentiation of *Vsig4*^+^ lining from *Aqp1*^+^ macrophages, we scored genes with differential expression and high ratios of unspliced transcripts along the pseudotime trajectory. This identified *Skil*, *Tal1*, *Nr4a3*, *Zmiz1* and *Z[x3* as potential transcriptional regulators of lining macrophage maturation (Figure 4D-E). At 3 weeks and in adult joints, an additional trajectory was predicted according to which *Ccr2*^+^ infiltrating cells produce *MHCII*^+^, *Retnla^+^* and *Cxcl9*^+^ IFN-activated sublining macrophages (Figure 4C), in line with our fate mapping results (Figure 3). Importantly, however, a direct trajectory from *MHCII*^+^ sublining to lining macrophages was not predicted, consistent with the limited BM contribution we observed in experimental fate mapping. Reassuringly, we obtained similar predictions about developmental trajectories with 2 alternative algorithms, Monocle 2 and diffusion maps (Figure S4A).

Next, we wanted to assess if the postnatal expansion and differentiation of lining macrophages is facilitated by proliferation, and if so, which subsets undergo proliferation. To address this experimentally, we used a genetic mouse model in which proliferating cells can be labelled owing to expression of *Mki67* (*Mki67*^CreERT2^:Rosa26^lsl-tdT)32^. Macrophages with transcriptional signatures of proliferating cells are most abundant in newborns and 1-week-old mice (Figure 2). We therefore induced labelling in *Mki67*^CreERT2^:Rosa26^lsl-tdT^ mice within the first week of life. Using this approach, approximately 35% of the VSIG4^+^ lining macrophages were labelled in adult joints (Figure 4F), indicating they originated from cells proliferating at the newborn stage. Of note, we also observed a similar degree of labelling in MHCII^-^ sublining macrophages and near-complete labelling of MHCII^+^ sublining macrophages. However, approximately half of the adult blood monocytes were also labelled, which suggests that this approach also labels long-lived BM progenitors that undergo proliferation in newborns. Nonetheless, we have demonstrated conclusively that monocytes only contribute minimally to lining macrophages. Their labelling in *Mki67*^CreERT2^:Rosa26^lsl-tdT^ mice thus cannot result from labelled monocytes. Instead, these data further support the notion that lining macrophages expand postnatally through proliferation of developmentally restricted precursors and/or pre-existing macrophages.

To better understand which population(s) undergo proliferative expansion to produce lining macrophages, we again leveraged our scRNAsequencing dataset. Neither of the two proliferating clusters does express *Vsig4*, and conversely, terminally differentiated *Vsig4*^+^ lining macrophages do not show a proliferative transcriptomic signature (Figure 2D, 2G). In contrast, many of the proliferating M-phase *Mki67*^+^ and S-phase *Mcm3*^+^ macrophages display high levels of *Aqp1*, especially in 1 week-old and 3-weeks-old joints (Figure S4B). Moreover, in newborn to 3-weeks-old mice, many of the proliferative *Mki67*^+^ and *Mcm3*^+^ macrophages were also enriched in a gene set that is characteristic of the *Aqp1*^+^ population (Figure 4G). This gene set comprises the genes most differentially expressed by adult *Aqp1*^+^ macrophages^1^. The proliferating macrophages enriched in this signature may thus represent *Aqp1*^+^ macrophages undergoing proliferation. Intriguingly, the gene signature of *Aqp1*^+^ macrophages is also enriched in *Vsig4*^+^ lining macrophages, further indicating a developmental relationship with *Aqp1*^+^ macrophages. Together with the differentiation trajectory inference, these analyses hence indicate that proliferating *Aqp1*^+^ macrophages provide a source for lining macrophages during postnatal development. In summary, lining macrophages differentiate and expand postnatally via proliferation, likely through *Aqp1*^+^ intermediates.

### Lining macrophage development depends on CSF1 and TGFβ signalling and involves mechanosensing through PIEZO1

Having established the developmental kinetics and cellular sources of synovial lining macrophages, we lastly investigated which signals support lining macrophage development. Macrophages reside and function within tissue niches that fulfil three main roles: (1) they provide trophic factors, (2) physical anchorage to macrophages and (3) imprint them through tissue-specific cues that drive differentiation programs, which are superimposed onto the core macrophage signature. In return, this enables macrophages to support the proper functioning of the tissue^17^. Lining macrophages are the outermost layer of the synovium, facing the synovial cavity and engaging in direct cellular contact only with lining fibroblasts. In contrast to macrophages, fibroblasts already form a continuous lining in neonatal joints (Figure S1A).

We therefore focused our efforts to delineate mechanisms underlying lining macrophage development on fibroblast-derived signals and tissue-specific factors. We first applied an unbiased approach to identify candidate cellular interactions by performing MultiNicheNet analysis on our developmental single cell RNAsequencing time course. This allowed us to interrogate potential signal and receptor pairs as well as downstream pathways. We compared interactions of fibroblasts with lining macrophages predicted as enriched in newborn and adult joints (Figure 5A). In newborn and 1-week-old joints (Figure 5A, S5A), predicted key signalling factors included the cytokines *Ccl12*, *Ccl3*, *Ccl6* as well as fibroblast-derived *Igf2*, to which macrophages may respond predominantly via CCR1 and Integrin alpha 6 (*Itga6*)-dependent pathways. Cellular interactions become more complex in 3-weeks-old and adult joints (Figure 5A, S5A). Our analyses predicted that in adult joints, lining macrophages respond to *TgS2* produced by lining fibroblasts, as well as *Csf1*, *Ang*, *Lama4*, and *Ccn1/2* from both lining and sublining fibroblasts. Sublining fibroblasts were further predicted to provide *Fgf2* to lining macrophages. Lining macrophages express transcripts for a range of receptors that would equip them to respond to these cues, including *TgSr1*, *Csf1r*, *Egfr* and *Fgfr1*. Predicted genes induced downstream of these pathways include *Fgf2* and *Itgav*, which may facilitate interactions with the extracellular matrix (ECM), immune surveillance and tissue maintenance. Finally, since they form an epithelial-like barrier via tight junctions^1^, we also interrogated cellular interactions within the lining macrophage population. Indeed, we identified *F11r* as a potential key player mediating cell-cell interactions between individual lining macrophages, the gene encoding junctional adhesion molecule A (Figure 5A).

**Figure 5.**
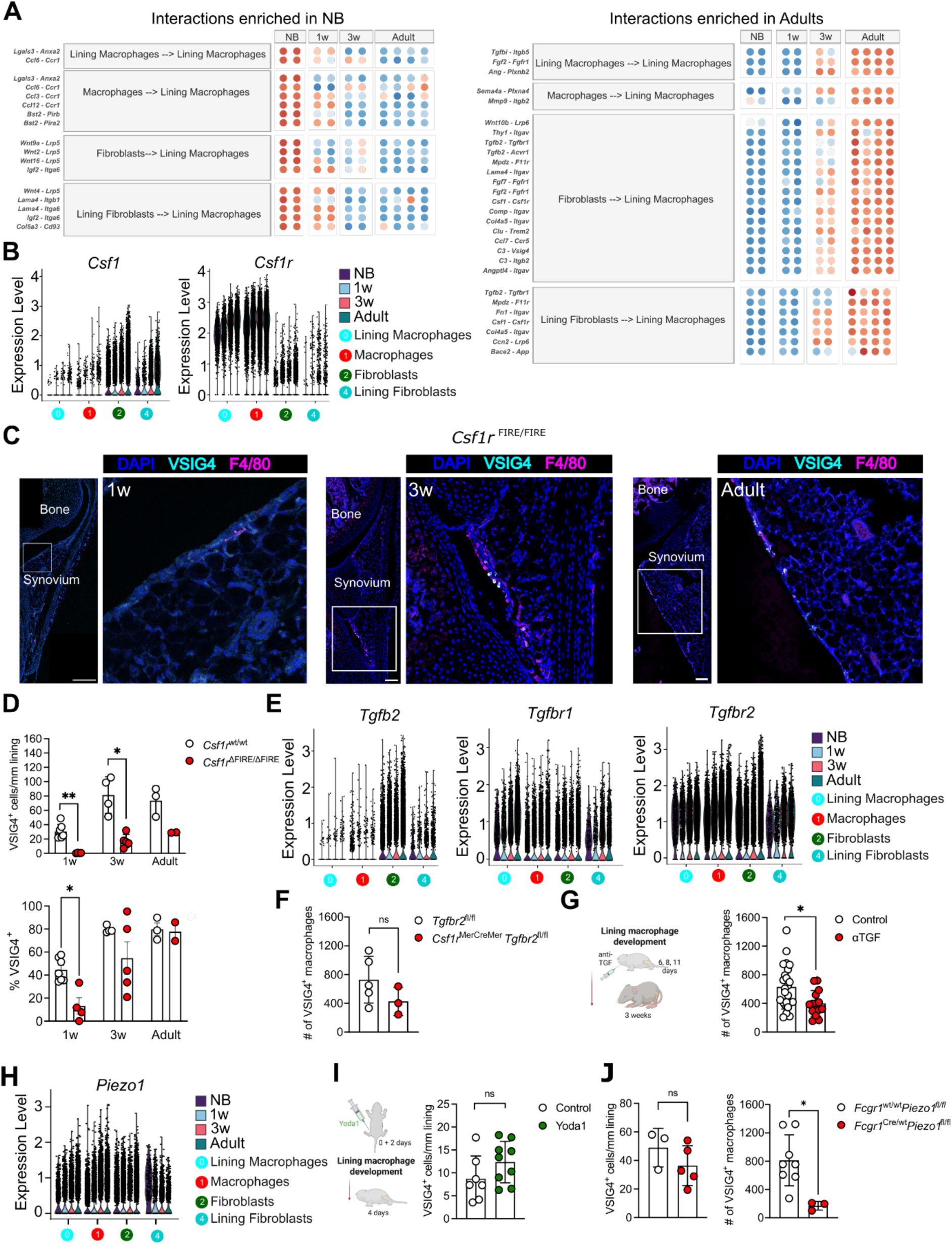
Lining macrophage development depends on CSF1 and TGFβ and involves mechanosensing through PIEZO1. **(A)** Computational inference of candidate signals instructing lining macrophage development. MultiNicheNet analysis of scRNAseq data. Bubble plots show the top pathways predicted to be differentially active in the newborn (NB) and adult stage. **(B)** Expression of *Csf1* and *Csf1r* by synovial macrophages and fibroblasts during postnatal development. **(C, D)** Synovial macrophage development in *Csf1r*^ΔFIRE/ΔFIRE^ mice. **(C)** Representative immunofluorescence images of synovial tissue of 1 week-old, 3 weeks-old and adult mice, stained for DAPI (blue), VSIG4 (cyan), and F4/80 (magenta). **(D)** Quantification of density (top) and frequency of VSIG4^+^ cells within lining macrophages (bottom) in the synovial lining of *Csf1r*^ΔFIRE/ΔFIRE^ mice compared to wild-type controls. **(E)** Expression of *Tgfb2* and *Tgfbr1* by synovial macrophages and fibroblasts during postnatal development. **(F, G)** Impact of TGFβ signalling on synovial macrophage development. TGFβ signalling was blocked in pups using antibodies or a genetic approach, and the abundance of VSIG4^+^ lining macrophages was determined at 3 weeks using flow cytometry. **(F)** *Csf1r*^MerCreMer^ *Tgfbr2*^fl/fl^ or Cre-negative *Tgfbr2*^fl/fl^ control pups were treated tamoxifen at 2 days of age and analysed at 3 weeks of age by flow cytometry. **(G)** Pups were treated with anti-TGF antibodies (clone 1D11) during postnatal development. Mice received 100µg antibody or PBS (Control) at 6, 8 and 11 days of age. **(H)** Expression of *Piezo1* by synovial macrophages and fibroblasts during postnatal development. **(I, J)** Synovial macrophage development in mice with loss or gain of function in PIEZO1 signalling. **(I)** PIEZO1 signalling was activated in pups using the specific agonist Yoda1. Pups received 70 ng per gram of body weight of the agonist or solvent (control) subcutaneously on the date of birth and at 2 days of age. The density of VSIG4^+^ macrophages was then assessed in 4-days-old pups by confocal imaging. **(J)** 3 weeks-old macrophage-specific *Piezo1* knockout mice (*Fcgr1*^Cre^:*Piezo1*^fl/fl^) were analysed by confocal imaging (left) and flow cytometry (right). Scale bars: 50 µm. For imaging analyses, each dot represents an individual mouse, with quantification based on 2 to 3 sections per mouse. For flow cytometry, each dot represents an individual mouse. Data are shown as mean ± SD. Statistical significance was determined using the Mann–Whitney test for pairwise comparisons. *p<0.05, **p<0.01, ***p<0.001.

We then experimentally tested *in vivo* the involvement of signals identified in this unbiased computational approach in lining macrophage development and/or maintenance. We focussed these analyses on FGF2, CSF-1 and TGF-β. Despite robust expression of *Fgf2* by sublining fibroblasts and *Fgfr1* by lining macrophages (Figure S5B), the density of VSIG4^+^ lining macrophages was unaffected in *Fgf2* knockout mice at 3 weeks of age (Figure S5C), indicating that FGF2 signals are not essential to the macrophage lining layer. CSF1 is a key factor required for the differentiation, proliferation, and survival of many macrophage populations. *Csf1* and *Csf1r* are highly expressed by (lining) fibroblasts and (lining) macrophages respectively (Figure 5B). To investigate if synovial lining macrophage development depends on CSF1, we performed a time course analysis of mice in which macrophages have reduced levels of CSF1R (in a tissue-specific manner) owing to the deletion of a super-enhancer region termed fms-intronic regulatory element (“FIRE”)^33^. Compared with *Csfr1*-proficient (*Csf1r*^Wt/Wt^) controls, the density of VSIG4^+^ lining macrophages was strongly reduced in the joints of *Csf1r*^ΔFIRE/ΔFIRE^ mice at all stages analysed (Figure 5C, 5D). The postnatal development of synovial lining macrophages thus critically depends on CSF1 signalling. Curiously, however, the few macrophages that localised to the lining of adult *Csf1r*^ΔFIRE/ΔFIRE^ mice did express VSIG4. This indicates that those remaining in the lining in the absence of CSF1 signalling may be appropriately imprinted by adulthood (Figure 5D).

Next, we investigated TGFβ as a candidate factor involved in lining macrophage development. Synovial fibroblasts express *TgS2* at increasing levels during development, whereas (lining) macrophages express transcript for its receptors *TgSr1* and *TgSr2* in a similar pattern (Figure 5E). Of note, *TgSr1* and *TgSr2* are also expressed by synovial fibroblasts, suggesting that they too may respond to TGFβ. The effects of TGFβ on synovial macrophage development may thus be direct, indirect through effects on fibroblasts, or a combination of macrophage-intrinsic and -extrinsic effects. To address macrophage-intrinsic effects, we used macrophage-specific conditional knockout mice for *TgSr2* (*Csf1r*^MerCreMer^:*TgSr2*^fl^), and induced ablation of *TgSr2* by tamoxifen administration in 2-day-old neonates to prevent TGFβ action from the early stages of lining maturation onwards. By 3 weeks of age, this caused a trend towards lower lining macrophage density (Figure 5F). To address a role for TGFβ signalling more globally, we also treated 1-week-old pups with depleting antibodies against TGF and analysed the effects when mice were 11 days old, reasoning that this would account for the most dynamic window of lining macrophage expansion. This resulted in an ∼1.5-fold reduction in lining macrophage density compared to vehicle-treated controls (Figure 5G).

Finally, we turned our attention to niche-specific factors that could mediate lining macrophage specification. Synovial joints are subjected to high mechanical pressure upon loading and shear stress due to the movement of the synovial fluid^34^. The developing joint experiences an increase in loading as walking develops, which is required for cartilage and tendon maturation^34–36^. Specifically, mechanical loading by walking generates cyclic hydrostatic forces, where bone compression is transduced to fluid pressure^37–39^. In mice, the onset of walking occurs during the first three weeks of life^40^. This coincides with the maturation window of the synovial macrophage lining that we have identified here and may thus be an instructing signal. To test this, we first made use of an optogenetic approach. *Acta1-* rtTA:tetO*-*Cre:*ChR2-V5*^fl^ mice, light-sensitive Channelrhodopsin-2 (ChR2) is expressed in skeletal muscles allowing for controlled muscle contraction using blue (455nm) light^41^. In this system, we induced short, daily bouts of unlilateral hind leg muscle contractions via blue light exposure in the immediate postnatal period (i.e., first 4 days postnatally and one day recovery before euthanasia), when pups are normally still largely immobile. We reasoned that if the onset of locomotion was indeed a driving force of lining macrophage development, then it should be accelerated following optogenetic muscle stimulation in neonates. This approach resulted in an increased density of VSIG4^+^ and total (F4/80^+^) macrophages in the exposed leg of some 5 days-old pups, but others showed the opposite trend (Figure S5D). To further investigate a role for mechanical loading in lining macrophage development, we then focused on mechanosensing. Macrophages can respond to mechanical stimulation^42–46^ through several mechanosensors^47^, but cyclic hydrostatic pressure has primarily been linked to the mechanosensitive ion channel PIEZO1^43,46^. Lining macrophages express high levels of *Piezo1* (Figure 5H), but not other mechanoreceptors (Figure S5E). This includes the first lining macrophages found at birth. Akin to what we observed following induced muscle contractions, neonatal activation of PIEZO1 through administration of its specific agonist Yoda1 resulted in an increased density of lining macrophages in some, but not all treated pups (Figure 5I). Lastly, we generated conditional knockout mice in which all (CD64^+^) macrophages are *Piezo1*-deficient (F*cgr1*^Cre^:*Piezo1*^fl^) mice. These showed a reduction in lining macrophage density at 3 weeks of age (Figure 5J), suggesting a possible involvement of PIEZO1-mediated mechanosensing in macrophage lining development. A similar trend was also apparent in *LysM*^Cre^:*Piezo1*^fl^:*Piezo2*^fl^ mice^48^ (Figure S5F).

Collectively, we demonstrated here that the formation of the macrophage lining is dependent on CSF1 and TGFβ signalling and may initially occur under the influence of cyclic mechanical pressure sensed through PIEZO1.

## Discussion

In this study, we comprehensively profiled joint macrophage development, focusing on those in the synovial lining. We found that in newborn mice and human foetuses, the lining is immature and only sparsely populated by macrophages. In mice, lining macrophages are established during the first weeks of life independently of monocytes. Instead, lining macrophages originate predominantly from foetal-restricted progenitors and expand through proliferation of macrophages present at birth, which may involve Aqp1^+^ intermediates. This process is CSF1- and TGFβ-dependent and further involves mechanosensing through PIEZO1. Our work identifies the early postnatal period as critical for synovial lining macrophage development.

### The synovial lining macrophage layer is formed in the first weeks of life

It is increasingly appreciated that environmental influences during development shape tissue-resident immune cell responses and influence disease trajectories, often long before the onset of clinical symptoms. This is also the case for rheumatic diseases^49,50^. Synovial lining macrophages facilitate resolution of joint inflammation^1,3^, but also contribute to early influx of inflammatory cells^4^. In many other tissues, macrophages are long-lived and established in early life, making them susceptible to environmental imprinting or “programming” by early life events^51^. Similar considerations may apply to synovial lining macrophages. We thus comprehensively delineated their developmental kinetics and origins and investigated the factors involved in this process. We found that the neonatal murine and foetal human synovial lining macrophage layer are incomplete. In mice we describe that the synovial lining macrophage layer is immature at birth, and forms within three weeks of birth, with further maturation occurring until adulthood. Whilst we identified the first weeks of life as the maturation window in mice, we cannot assess later foetal (> 19 weeks EGA) and early postnatal human tissue due to availability. However, we did observe an increase in macrophage density in the developing lining area towards the later foetal stages analysed, suggestive of ongoing recruitment to the lining, similar to newborn mice. Postnatal macrophage maturation is consistent with other organs, where macrophage populations are undergoing maturation alongside their tissue niches^52,53^. Indeed, lining macrophages not only increase in density during the first three weeks of life, they also upregulate *Vsig4* expression and acquire an overall pro-inflammatory gene expression profile. In the healthy adult joint, lining macrophages form an epithelial-like barrier, closely engaging with each other through tight junctions facilitated by JAM1, ZO-1 and Claudin 5^1^. We report that lining macrophages acquire this characteristic feature between birth and three weeks of age, when they transition from an initially scattered distribution to a closed layer without gaps, concomitant with upregulation of *F11r* (encoding JAM1). Furthermore, within this 3-week window, CX3CR1 expression becomes increasingly restricted to lining macrophages, reminiscent of other tissues. Indeed, all developing tissues are initially seeded with CX3CR1^+^ macrophages^26,54^, whereas only certain adult populations retain CX3CR1. These are often long-lived, self-renewing macrophages, such as microglia^55^, arterial wall macrophages^56^ and nerve-associated dermal macrophages^57^.

### Other synovial macrophages are highly dynamic during postnatal development

In addition to those in the lining, murine and human joints host a range of other macrophage populations. These differ in gene expression profiles, localisation, and (potential) functions^1,3^. We confirmed all macrophage populations that had been identified in full ankle tissue^1^ in our single cell transcriptomic dataset from microdissected knee joints. Interestingly, we also identified populations that have not previously been described. These vary substantially in abundance at distinct life stages. *Clec4n*^+^ “developing” macrophages are most abundant in the neonatal synovium and gradually decrease, although not fully disappear from adult joints. It is unclear how this contraction occurs. Gene ontology (GO) analysis suggested an enrichment in translation-related processes through ribosomal proteins (*Rpl34*, *Rpl6*, *Rpl32*, *Rps11*) and in endocytosis/phagocytosis via Fc receptors (*Fcgr3*, *Fcer1g*, *Fcgr2b*) and vesicle-related genes (*Cd63*, *Ap2m1*, *Cyba*, *Ap2m1*). These macrophages also express the cytokine transcripts *Ccl12*, *Ccl7* and *Ccl2*, which could implicate them in monocyte recruitment and blood vessel formation during synovial tissue maturation. *Egfr*^+^ macrophages follow a similar developmental pattern to lining macrophages. They appear to be primed for differentiation, as indicated by high levels of transcriptional regulators such as *Jun*, *Junb*, *Maf*, *Cited2*, and *Klf4*. Of note, although both *Clec4n*^+^ and *Egfr*^+^ macrophages highly express *Pde4c* in 1- and 3-week-old mice, inhibition of phosphodiesterase 4 activity with rolipram did not affect synovial tissue development (Figure S5G). It is important to note that we cannot rule out contamination with circulating cells, especially at the younger life stages, where microdissection is more challenging. Indeed, whilst we found *Ly6c*^+^ monocytes and *Ccr2*^+^ macrophages in all samples, they are relatively more abundant in neonates, which could be due to more contamination from the circulation. However, *Clec4d*^+^ “activated” macrophages express high levels of *Timd4*, which is associated with tissue residency. *Cxcl9*^+^ “IFN-activated” macrophages, on the other hand, are restricted to the adult joints, and their characteristic IFN-signature suggests they may be involved in immune surveillance and inflammatory responses within the mature synovial tissue. Although beyond our current focus on lining macrophages, investigating the potential stage-specific roles of these macrophage populations during synovial tissue maturation and maintenance is an exciting avenue for future studies, as is their relevance to joint inflammation, resolution and pathological remodelling.

### Ontogeny of synovial lining macrophages

Using a range of complementary genetic fate mappers, we show that synovial lining macrophages originating from foetal-restricted progenitors colonising the synovial lining during the first 3 postnatal weeks. This process involves proliferation but is independent of monocyte recruitment. As animals grow into adulthood (6-8 weeks), the lining undergoes further expansion. In this period, a small pool (approximately 10%) of GMP-derived monocytes does integrate into the lining. Overall monocyte contribution is then maintained at a similar level until 6 months of age. This is consistent with two recent reports showing limited monocyte labelling in adult *Ms4a3*^Cre^ fate mappers^12,58^, but provides higher temporal resolution. Intriguingly, pulse-chase labelling of HSCs indicates a similar frequency of turnover in adult mice, suggesting that the few monocyte-derived lining macrophages might get continuously replenished from the bone marrow. Moreover, monocytes can also originate from bone marrow MDPs, and these can produce specific types of macrophages^18,19^. We demonstrated here, for the first time, that synovial lining macrophages do not originate from MDP-derived monocytes. Compared to macrophages in other tissues, this degree of monocyte dependence is very low, second only to microglia and comparable to liver Kupffer cells and epidermal Langerhans cells^14,29,59,60^.

Instead of monocyte recruitment, our data indicated that postnatal expansion and adult maintenance of lining macrophages involves *Aqp1*^+^ intermediates from the sublining. A population of CX3CR1^-^ sublining macrophages has previously been reported as a source of lining macrophage replenishment in adult joints^1^, CX3CR1^-^ primarily comprise *MHCII*^+^, *Retnla^+^*, and *Aqp1*^+^ macrophages^1^, but VSIG4^+^ lining macrophages do not have a history of *Retnla* expression, making *Aqp1*^+^ macrophages the likely source. In line with this, Schonfeldova and colleagues have now characterised AQP1^+^ macrophages associated with the meniscus area, which are also largely monocyte-independent and maintained through local proliferation^31^. Since lining macrophages are long-lived, supplementation from the AQP1^+^ pool may be a way of fast-tracking replenishment of the lining when homeostasis is perturbed.

Taken together, our data and these studies suggest that AQP1^+^ macrophages are a source of lining macrophages during postnatal maturation and in healthy and inflamed adult joints.

### Factors involved in lining macrophage development

In contrast to macrophages, lining fibroblasts already appear fully established at birth. Nonetheless, they undergo substantial changes during postnatal development, displaying an overall shift from an inflammatory tissue-remodelling to an anti-inflammatory signature. Macrophages integrate into this structure later, suggesting a coordinated crosstalk between fibroblasts and macrophages (and their progenitors) during development^61^. Indeed, in an *in vitro* co-culture system, synovial fibroblasts spontaneously incorporate monocytes into a lining layer^62^, and our transcriptomic data predict that fibroblasts are the main signal-sending cells in the synovium.

Lining fibroblasts express *Csf1*, which we identify here as a key trophic factor for lining macrophages during development. This confirms previous reports in osteopetrotic (op/op) mice, which lack the *Csf1* gene^63^. Of note, we observed that the few macrophages remaining in the lining of *Csf1r*^ΔFIRE/ΔFIRE^ mice are VSIG4^+^, indicating that CSF1 signalling is primarily needed for survival within and/or recruitment to the lining, but not imprinting of the niche-specific identity, and reinforces the central role of CSF1 in macrophage biology, in keeping with a “division of labour” within the lining macrophage niche. We further found that TGFβ signalling contributes to lining macrophage development within the maturation window. Similar requirements for TGFβ have been described for the development of microglia^64^, Langerhans cells^65^, alveolar macrophages^66^ and osteoclast differentiation^67^. Our transcriptomic data suggest that synovial fibroblasts may be both, producing and sensing TGFβ. Indeed, we observed a stronger reduction in lining macrophage density when TGFβ was blocked using antibodies, rather than a macrophage-specific knockout for *TgSr2*, indicating that the effects of TGFβ on lining macrophage development are at least in part also in a non-cell-autonomous manner. Moreover, TGFβ may also signal through TGFBR1 in macrophages, which we cannot account for in our genetic model.

Based on the observation that the maturation of the macrophage lining coincides with the onset of walking in mice, we hypothesised that mechanical loading may function as an instructing signal in this process. In agreement with this hypothesis, we observed a reduction in lining macrophage density at 3 weeks of age in the joints of mice with a macrophage-specific knockout for the mechanosensory *Piezo1* (*Fcgr1*^Cre^:*Piezo1*^fl^). A similar trend was also observed in *LysM*^Cre^:*Piezo1*^fl^:*Piezo2*^fl^ mice. Conversely, neonatal pharmacological activation of PIEZO1 or optogenetically induced muscle contractions both resulted in an increased density of lining macrophages in some, but not reliably in all pups. This might reflect background levels of mechanical loading in our loss- and gain-of function approaches. Moreover, different types of mechanical forces and mechanosensitive pathways could contribute to lining macrophage specification during development, and even at adult stages. For example, static forces likely also apply during postnatal development owing to tissue growth and elongation. It is conceivable therefore that different types of mechanical forces and pathways of mechanosensing integrate to mediate lining maturation during postnatal development.

In conclusion, our work identifies the first weeks of life as the critical window and CSF1 and TGFβ as key signals for the development of synovial lining macrophages. This may have implications for joint health and disease across the lifespan, including juvenile arthritis and how disease susceptibility is shaped by the early life environment.

## Acknowledgements

We thank the IRR Flow Cytometry & Cell Sorting Facility, the IRR Single cell multi-omics facility, the IRR Imaging facility and the University of Edinburgh Bioresearch & Veterinary Services for their invaluable services and experimental support. Sequencing of single cell libraries was carried out by Edinburgh Genomics and the Edinburgh University Genetics Core. The Edinburgh Compute and Data Facility (ECDF) are acknowledged for providing essential resources (http://www.ecdf.ed.ac.uk/). We thank all donors who generously contributed human foetal tissue, and Richard Anderson and Norma Forson for providing these samples. We also thank Clare Pridans for *Csf1r*^ΔFIRE^ mice. Finally, we thank Florent Ginhoux, Hongkui Zeng, and Steffen Jung for providing us with the sequences for the Cre, tdTomato, and GFP DNA constructs.

## Author contributions

Conceptualization: MMP, BMD, RG

Data curation: MMP, BMD, RG, AA, GS, GD, JK, SU, MLK

Formal analysis: MMP, BMD, RG, AA, GS, GD, JK, Funding acquisition: RG

Investigation: MMP, BMD, RG, AA, GS, GD, JK, SB, KP, JZ, EE, CM, NM, MK, OBB, SU, AG, SI Methodology: CCB, EE, BS, KZ, TS, LM, JB, GN, SK, CM, AC, NM, MK, OBB, SU, AG, SI Resources: CR, IU, MKS, TV, RR, BWM, AC, LB, NM, MK, DV

Visualization: MMP, BMD, RG Writing, original draft: MMP, BMD, RG

Writing, review & editing: All authors critically revised the manuscript and agreed to the final submission.

## Resource availability

All data related to this study are available either in the paper or in the Supplementary Materials. Requests for further information and resources should be directed to and will be fulfilled by the lead contact, Rebecca Gentek.

## Materials and Methods

### Human samples

Human fetal tissues were obtained with written informed consent immediately after medical elective termination of pregnancy. The study was approved by the Lothian Research Ethics Committee (ref 08/S1101/1), Scotland, UK.

### Animal studies

All animal procedures were approved by the UK Home Office and performed under project license PP1871024 according to the AWERB institutional guidelines. Mice of both sexes were used in experiments and randomized between cages to minimize cage effects. Transgenic mouse lines used in this study are listed in Supplementary Table 1. Mice were housed under standard specific pathogen-free (SPF) conditions, with unrestricted access to food and water, and maintained on a 12-hour light/dark cycle. Mice were humanely killed using a Schedule 1 method, in accordance with UK Home Office guidelines. No statistical methods were employed to predetermine sample sizes for the experiments.

### Tamoxifen administration and cell labelling strategies

Newborn mice were treated with tamoxifen (Sigma-Aldrich) for labeling of CX3CR1⁺ cells (*Cx3cr1*^CreERT2^:Rosa26^lsl-tdRFP^) or proliferating cells (*Mki67*^CreERT2^:Rosa26^lsl-tdT^), or for macrophage-specific deletion of *TgSr2* (*Csf1r*^MerCreMer^:*TgSr2*^fl^). Mice were administered a single dose of tamoxifen intraperitoneally (IP) at 25 µg per gram of body weight in corn oil. For adult cell labeling, *Cxcr4*^CreERT2^:Rosa26^lsl-tdT^ and *Kit*^MerCreMer^:Rosa26^lsl-eYFP^ mice were treated with tamoxifen dissolved in corn oil. Tamoxifen was delivered by oral gavage at 0.12 mg per gram of body weight per day for three consecutive days. For all treatments, tamoxifen solutions were sonicated prior to administration.

Fetal fate mapping was performed using *Cdh5*^CreERT2^:Rosa26^lsl-tdT^:*Cx3cr1*^gfp^ and *Cx3cr1*^CreERT2^:Rosa26^lsl-tdRFP^ mice. Female mice were timed-mated, and the presence of vaginal plugs the morning after mating was considered embryonic day 0.5 (E0.5). To induce reporter recombination in the developing offspring, a single dose of 4-hydroxytamoxifen (4OHT, 1.2 mg) (Sigma-Aldrich) was administered to pregnant females via IP injection at E7.5, E9.5 or E10.5 as indicated. To mitigate the adverse effects of 4OHT on pregnancy, progesterone (0.6mg) (Sigma-Aldrich) was co-administered. In cases where dams were unable to deliver naturally, pups were delivered via cesarean section and cross-fostered by lactating CD1 mice.

### Other *in vivo* treatments

Rolipram was dissolved in ethanol at 24.65mg/ml, diluted in PBS, and administered via IP injection at 0.5 µg per gram of body weight on postnatal days 3 (P) 3, P5 and P7 with analysis on P21. Anti-TGF depleting antibodies (clone 1D11) were diluted in PBS and injected at 100µg per injection on P6, P8, and P11, with analysis on P21. The Piezo1 agonist Yoda1 was dissolved in DMSO at 5mg/ml, diluted in PBS, and administered subcutaneously at 70 ng per gram of body weight on P0 and P2, with analysis on P4. In all experiments, PBS-injected littermates served as controls.

### Optogenetic induction of muscle contractions

*Acta*-rtTA; tetO-Cre; *ChR2-v5*^fl/fl^ male mice were crossed with *ChR2-v5*^fl/fl^ females and pregnant dams were given doxycycline chow during pregnancy. Cre^+^ pups were genotyped by optogenetic stimulation and confirmed using PCR. Newborn pups were laid on their right side and their left hind limb was unilaterally stimulated using blue light to induce unconstrained skeletal muscle contractions (50 cycles per bout; each cycle consisted of 70msec light on, 30msec light off for 1 second with 3 seconds rest between cycles). These contractions resulted in cyclic extension (when the light was on) and relaxation (during rest) of the hindlimb. This was repeated for four consecutive days (P0-P4), and pups were euthanized at 5 days of age. Hindlimbs were collected and fixed in cold 4% paraformaldehyde, then stored in sodium azide prior to cryosectioning.

### Tissue Digestion and Flow Cytometry

The knee synovial tissue block, encompassing the supra- and infrapatellar synovial lining, associated joint capsule, infrapatellar fat pad, patella, patellar tendon and quadriceps tendon, was micro-dissected from knee joints following a previously described protocol^11^. Briefly, the skin was removed from hind legs to expose muscles and the underlying synovial tissue block. Excess muscle tissue was trimmed from the femur, exposing the patella and quadriceps tendon. The quadriceps tendon was cut approximately 3-4 mm proximally to the patella and used to pull the synovial tissue block away from the joint. Whilst gently applying constant pressure, the edges of the joint capsule were cut, liberating the synovial tissue block, which was then removed by cutting along the tibial portion of the joint capsule. Isolated synovial tissue blocks were kept in RPMI 1640 medium (Gibco) + 2% Fetal bovine serum (FBS) until digest.

Single cell suspensions from the brain and synovial tissue block were obtained through mechanical dissociation and enzymatic digest in a solution containing 0.2 mg/ml DNaseI (Roche), 400 U/ml Collagenase I (Gibco), and 2 mg/ml Dispase (Roche) in RPMI 1640 medium (Gibco) + 2% FBS. Digests were conducted for 20-30 minutes at 37°C under continuous horizontal shaking (900rpm). To support mechanical dissociation and digestion, samples were pipetted up and down regularly. Blood samples collected in EDTA tubes underwent red blood cell (RBC) lysis for 3 minutes using home-made RBC lysis buffer (150 mM NH4Cl, 10 mM NaHCO3, 1.2 mM EDTA in H2O). All digested tissue samples were filtered (70 μm) before staining.

To block and stain for viability, cells were incubated with TruStain FcX (Biolegend) in rat- and mouse serum-supplemented FACS buffer and stained using the LIVE/DEAD Fixable Near-IR Dead Cell Stain Kit (Invitrogen). Subsequently, cells were extracellularly stained for 20-30 minutes with fluorochrome-conjugated antibodies (Supplementary Table 2) in FACS supplemented with 20% BD Horizon Brilliant Stain Buffer. Cells were either fixed or not with Antigenfix (DiaPath). Flow cytometry was performed on the BD 5-laser LSR Fortessa. FACSDiva software was used for data acquisition, and data were analyzed with FlowJo (LLC, V10.8.1). A representative gating strategy is shown in Figure S2A.

### Cell sorting and single cell RNA sequencing (scRNAseq)

Synovial cells were isolated and stained as described above followed by cell sorting through a 70-μm nozzle using the Fusion Q cell sorter. Prior to sorting, DAPI (4ʹ,6-diamidino-2-phenylindole) (Thermo Fisher) was added to the cell suspension to exclude dead cells. Sorted populations included non-neutrophil immune cells (DAPI^-^ CD45^+^ Ly6G^-^) and fibroblasts/endothelial cells (DAPI^-^ CD45^-^ CD31^+^ or Podoplanin^+^). Sorting was conducted based on the gating strategy shown in Figure S2A. Post-sort cell count showed that all sorted populations exhibited a viability exceeding 90%. Before loading cells sorted cells to the 10X platform, the immune cells were pooled with the sorted fibroblasts/endothelial cells at a ratio described in Supplementary Table 3.

### scRNAseq: Library preparation

Gene expression (GEX) and multiplex (CMO) libraries were generated using the Chromium Single Cell 3’ Reagent Kit V3.1 from 10X Genomics following the manufacturer’s instructions. Each sample loaded onto a 10X Genomics cartridge consisted of up to 60,000 cells. Different litters or sexes were pooled before sorting using the 3’ CellPlex Kit Set A, following the manufacturer’s instructions (10X Genomics). GEX and CMO libraries were purified using SPRIselect beads (Beckman Coulter). The libraries were quantified and assessed for size distribution on the Qubit 2.0 Fluorometer (Thermo Fisher) and the Agilent Bioanalyzer (Agilent Technologies) using the Qubit dsDNA HS assay Kit and the DNA HS Kit, respectively. Results were utilized to calculate the library molarity. Library preparation and quality control were conducted at the IRR Cell Sorting Facility. GEX libraries and CMO libraries were pooled together with PhiX Control v3 (∼1%) and sequenced on the NextSeq 2000 platform or the NovaSeq 6000 platform using the NextSeq 1000/2000 P3 Reagents (100 cycles) v3 Kit or the NovaSeq 6000 S2 Reagent (100 cycles) Kit v1.5 following a paired-end reading strategy. Sequencing of the single cell libraries was performed by Edinburgh Genomics and the Edinburgh Genetics Core. Reads were converted to the fastq format using the Cell Ranger mkfastq pipeline (10X Genomics, V7.1.0). Subsequently, raw gene expression matrices were generated and demultiplexed per sample using the Cell Ranger multi-pipeline (10X Genomics, V7.1.0). Before alignment, the STAR indexes for read alignment were built by combining the mouse reference genome (GRCm38) with the corresponding genomic sequence of Cre, GFP, tdTomato.

### scRNAseq analysis: Gene expression and pathways

All packages (version) utilized for the analysis and visualization of scRNAseq data are summarized in Supplementary Table 4. Individual raw gene expression matrices per sample were merged and analyzed using the Seurat package. The cell matrices underwent filtering, removing cell barcodes with < 200 features, >5101 features (95th percentile cutoff), >18801 expressed genes (95th percentile cutoff), or >7.5% of reads mapped to mitochondrial RNA. The remaining cells were normalized using the LogNormalize method, and the 3000 most variable genes were selected for principal component analysis (PCA) to reduce dimensionality and cluster all cell types. The selection of principal components was based on elbow and Jackstraw plots (typically 40–50). Harmony, with a theta value of 0, was applied to correct for different sample lanes. After quality filtering, we obtained ∼630 million unique transcripts from 60,929 cells, each with more than 200 genes, averaging 3,160 genes per cell. This included 13,050 (21.4%), 11,961 (19.6%), 13,876 (22.8%), and 22,042 (36.2%) cells from Newborns (NB), 1-week-old (w1), 3-week-old (w3), and adult mice, respectively.

Clusters were determined using the FindClusters function with a resolution between 0.2 and 2 and visualized using Uniform Manifold Approximation and Projection for Dimension Reduction (UMAP). Differential gene expression (DGE) analysis between clusters, conditions, and tdTomato^+^ vs. tdTomato^-^ cells was performed using the FindMarkers function. DEGs with an adjusted P-value < 0.05 were subjected to pathway analysis using the Gene Ontology database. Gene module scores were calculated in Seurat using the AddModuleScore function. A predefined gene list of all synovial macrophages subsets was derived from Culemann *et al*. ^1^. Cells with a higher score demonstrate greater enrichment for the given gene module. These scores were subsequently utilized for downstream visualization and clustering analyses.

Clusters were annotated to cell types based on the expression of marker genes. We revealed 19 distinct cell types based on key transcriptional markers (Figure 2 and Figure S2). Computationally, we isolated and subclustered macrophages and fibroblasts. Re-clustering, non-linear dimensionality reduction, and analysis were performed as described above.

### scRNAseq analysis: Cell-to-cell communication

Cell-cell communication was inferred using the MultiNicheNet package, which predicts ligand-receptor interactions and downstream signaling based on pathway information. The default MultiNicheNet pipeline was applied to compare each timepoint, enabling identification of timepoint-specific cell-cell communication. Analyses were performed using all cell clusters, with a focus on (lining) fibroblasts and (lining) macrophages as sender populations and lining macrophages as the receiver population.

### scRNAseq analysis: Developmental trajectories and RNA velocity

The R packages Monocle3^74^ and Slingshot^75^ were employed to investigate pseudotime trajectories within the macrophage compartment. For Monocle analysis, Seurat objects were transformed into cell_data_set objects. Monocle trajectory analysis utilized Seurat clustering information and UMAP choosing monocytes as the root. In the case of Slingshot, Seurat objects were transformed into SingleCellExperiment objects. Slingshot trajectory analysis was conducted using Seurat clustering information and diffusion map dimensionality reduction. Additionally, RNA velocity analysis was performed following a previously described method. Initially, loom files containing spliced and unspliced counts were generated using Velocyto^76^. These loom files were loaded into Scvelo to estimate and visualize RNA velocities according to the standard pipeline^77^. The partition-based graph abstraction (PAGA)^78^ was computed based on the RNA velocity graph.

### Tissue processing for imaging

NB, w1, w3, and adult mouse legs were fixed in 4% paraformaldehyde (PFA, Thermo Fisher) at 4°C for 1h, 1.5h, 3h, and overnight, respectively, followed by a PBS wash. Decalcification was performed on all legs except for newborn samples in 0.5 M EDTA at 4°C for 3 days (w1) or 10 days (w3 and adult). Following decalcification, tissues were dehydrated by incubation in 30% sucrose overnight at 4°C. Tissues were then equilibrated in a 1:1 mixture of 30% sucrose and optimal cutting temperature (OCT) compound (Leica) for 3–5 days prior to embedding in OCT. Frozen sections (16–18 μm thick) were cut (Superfrost Microscope Slides, epredia) using a Leica cryostat and stored at −20°C.

### Lightsheet microscopy

Knee samples from newborn mice were fixed overnight in Fix/Perm buffer (BD Bioscience) and subsequently washed in PBS. Immunostaining was carried out using anti-GFP AlexaFluor 488 (Biolegend, clone FM264G, 1:50 dilution) and anti-CD44 AlexaFluor 647 (Biolegend, clone IM7, 1:100 dilution) in PBS containing 0.2% Triton X-100 (Sigma-Aldrich) overnight at room temperature. Following staining, samples underwent three one-hour washes in PBS/0.2% Triton X-100. Dehydration was achieved by sequential incubation in ethanol at increasing concentrations (50%, 70%, and 100%), each for one hour at room temperature, followed by refractive index matching in ethyl cinnamate (Sigma-Aldrich) overnight at room temperature. Samples were imaged using a Luxendo MuVi SPIM lightsheet microscope equipped with 488nm and 643nm lasers with a 4.9μm beam expander for double-sided illumination, and bandpass filters of 497–554nm and 655–704nm for detection. Imaging was performed with a 10x detection lens with 10x internal magnification, acquiring overlapping tiles in four rotational stacks (90° increments). Image channels were aligned and the four perspectives were registered and merged using LuxBundle software (Luxendo/Bruker, v5.0.3) with manually selected landmarks to generate an isotropic 3D dataset with voxel dimensions of 1.30μm in x, y and z dimension. Image reconstruction, analysis, manual masking of the CD44-high parietal lining surface, and visualization were conducted using IMARIS software (Bitplane, v10.2.0).

### Immunofluorescence Imaging

Frozen slides were equilibrated to room temperature and subsequently incubated at 37°C overnight to improve tissue adherence. Tissue sections were permeabilized and blocked for 30 min at room temperature using 0.5% Triton X-100 and 1% bovine serum albumin (BSA) (Sigma-Aldrich) in PBS. Slides were then incubated with primary antibodies (listed in Supplementary Table 5) overnight at 4°C. Following primary antibody staining, tissues were incubated with Alexa Fluor 647 (AF647)- or AF488-conjugated secondary antibodies (listed in Supplementary Table 6) for 2 hours at room temperature. Lipid droplets were stained (1/1000, 1h) using BODIPY 493/503 (Thermo Fisher Scientific). Slides were mounted using fluoromount-G containing 1µg/mL DAPI (Thermo Fisher Scientific) prior to the addition of coverslips. Fluorescent images were acquired using a Leica TCS SP8 confocal microscope equipped with an Acousto-optical beam splitter. Z-stacks of images were captured. Blue diode (405nm), Argon (488nm), Diode (561nm, Hene (633nm).

### Imaging Analysis

Image analysis was performed the ImageJ software (Version 2.16.0). Maximum intensity z-stack projections were generated prior to analysis. Images were split into individual channels, and the relevant fluorescent channel was extracted for analysis. Positively stained cells for each marker were manually counted by at least two independent observers. For spatial analysis of F4/80+ cells, distances between individual F4/80+ cells were measured. The distance between two F4/80^+^ cells was defined as the shortest line connecting their respective peripheries. In cases where F4/80^+^ cells were directly adjacent, the distance was recorded as 0µm.

## Statistical analysis and reproducibility

All datasets were assumed to have a non-normal distribution. For comparisons between two groups defined by a single factor, the Mann-Whitney U test was applied. For comparisons among multiple groups, the Kruskal-Wallis test with Bunn’s corrections for multiple comparisons was used. Statistical significance was defined as p < 0.05. Statistical significance is denoted as follows: p < 0.05 (*), p < 0.01 (**), p < 0.001 (***), and p < 0.0001 (****). All statistical analyses and data visualizations were performed using GraphPad Prism (v10.1.2). In all figures, individual data points are displayed, and bars represent the mean ± SD.

---

**Figure S1.**
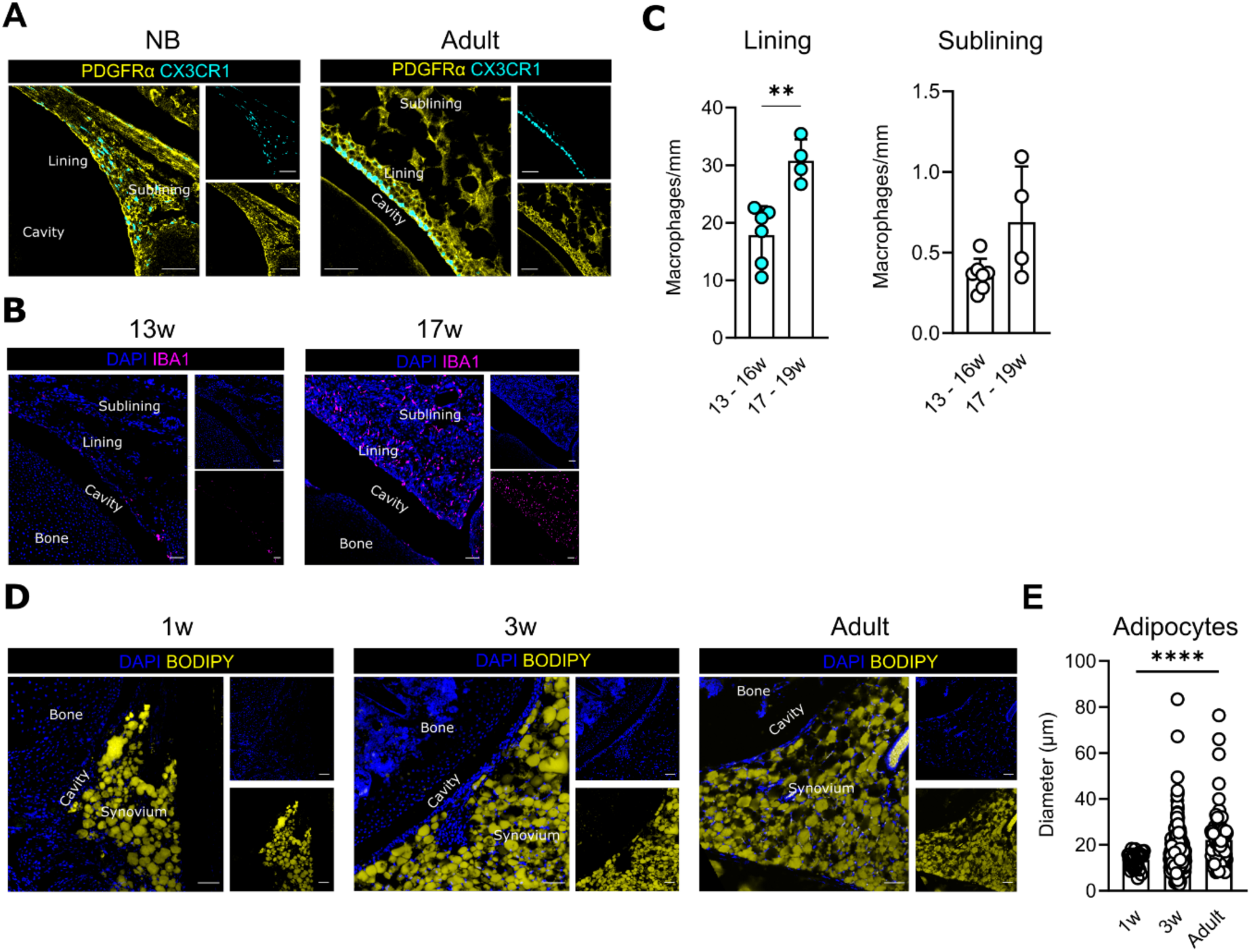
Developmental changes in synovial macrophages, fibroblasts and adipocytes. **(A)** Representative immunofluorescence images of newborn (NB) and adult mouse synovial tissue, showing PDGFRα⁺ fibroblasts (yellow) and CX3CR1-GFP⁺ cells (cyan). **(B)** Representative immunofluorescence images of human foetal synovium at 13 weeks and 17 weeks estimated gestational age (EGA), stained for DAPI (blue) and IBA1⁺ macrophages (magenta). **(C)** Quantification of IBA1⁺ macrophage density in the synovial lining and sublining of the human foetal synovium at 13 to 16 weeks and 17 to 19 weeks EGA. Each dot represents an individual foetus, with quantification based on 2 to 3 sections per foetus. **(D)** Representative immunofluorescence images showing adipocyte distribution in the synovium at the indicated stages, stained with DAPI (blue) and BODIPY (yellow). **(E)** Quantification of adipocyte diameter. Each data point represents a single adipocyte, and data were obtained from two individual mice per time point. Scale bars: 50 µm. Data are presented as mean ± SD. Statistical significance was determined using the Mann–Whitney test for pairwise comparisons or the Kruskal–Wallis test with Dunn’s correction for multiple comparisons, relative to NB. **p<0.01, ****p<0.0001.

**Figure S2.**
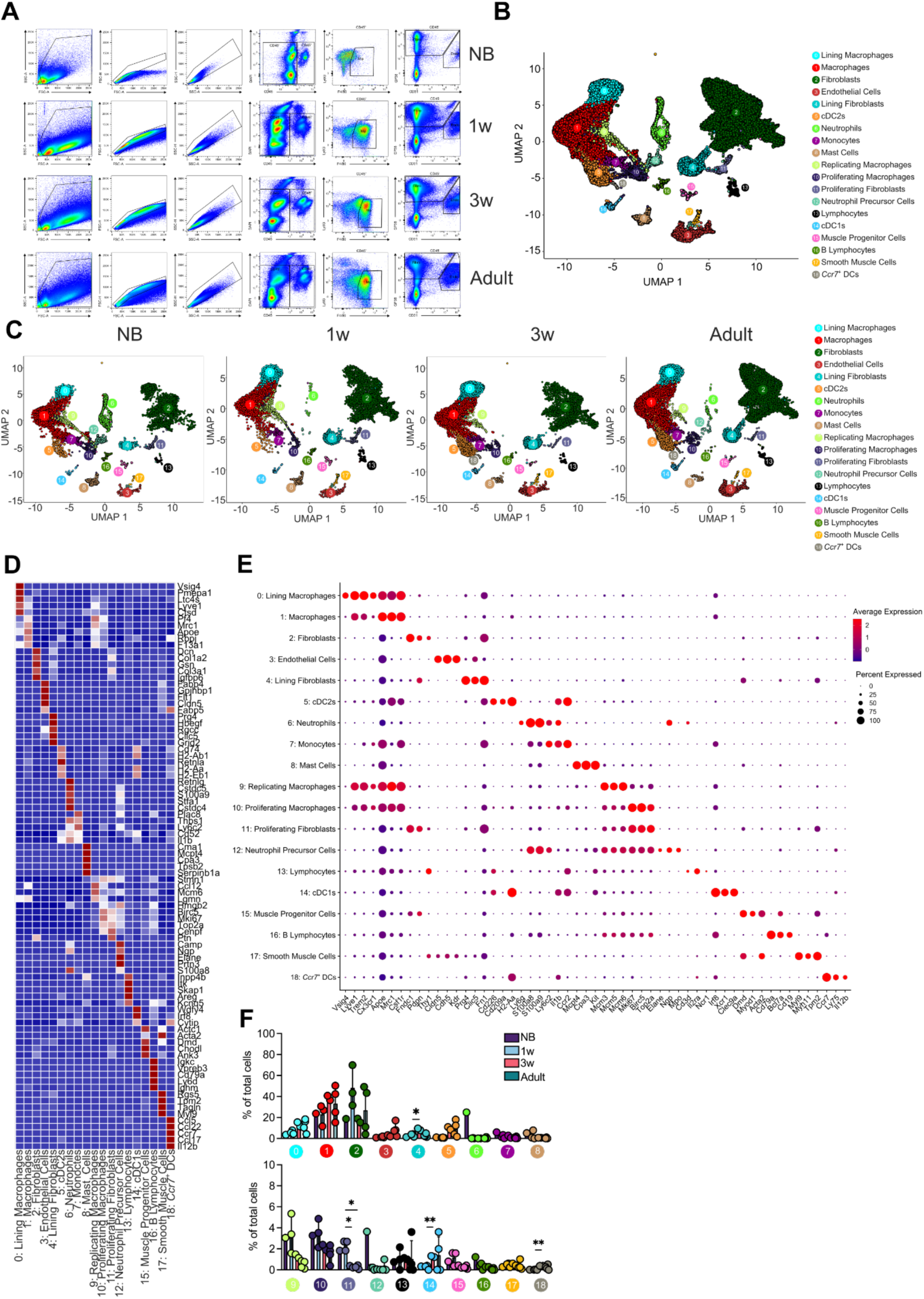
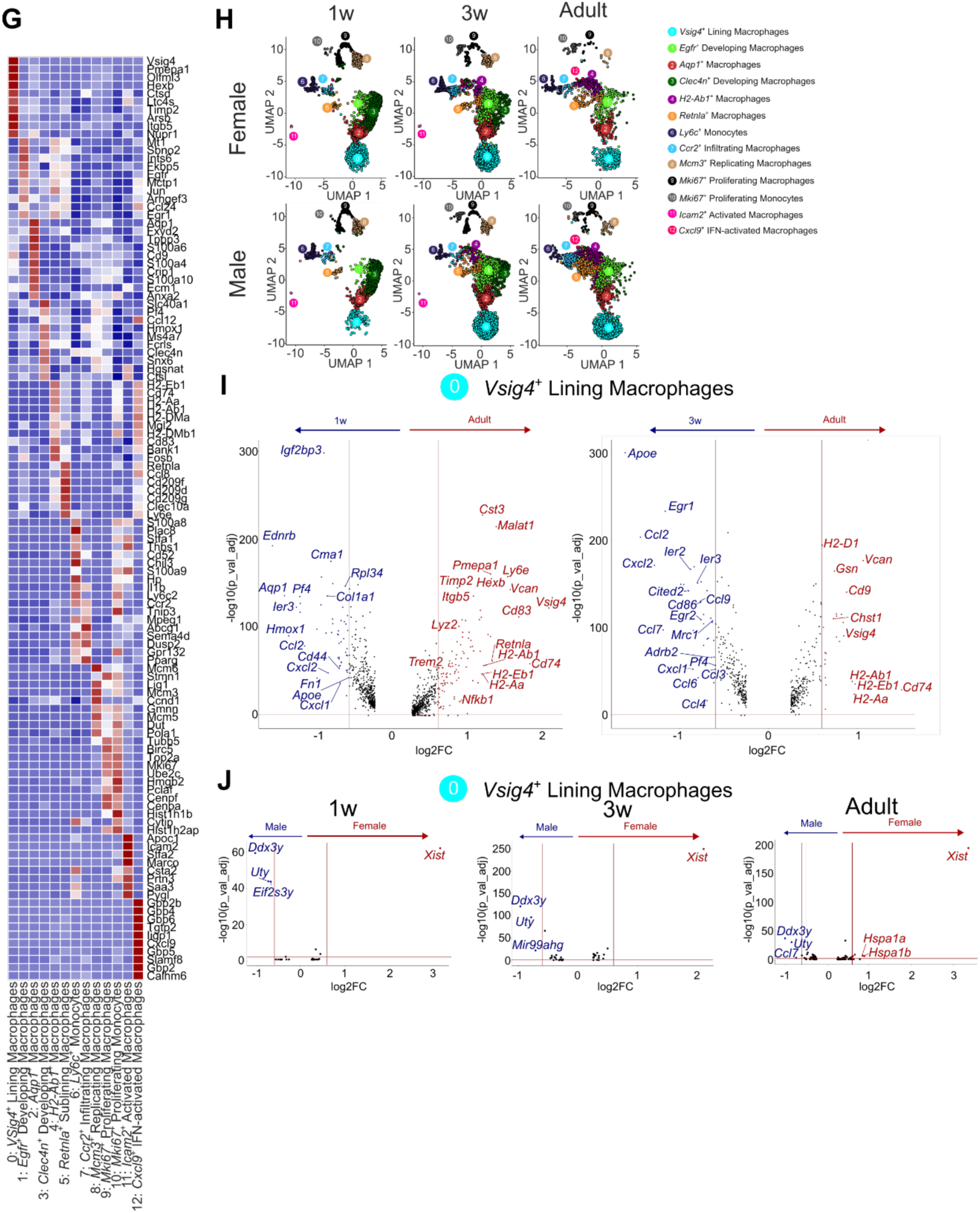
Flow cytometric identification and single-cell transcriptomic analysis of synovial cells. **(A)** Flow cytometry gating strategy used to sort macrophages and monocytes (live CD45⁺ F4/80⁺ Ly6G⁻), fibroblasts (live CD45⁻ GP38⁺), and endothelial cells (live CD45⁻ CD31⁺) from newborn (NB), 1-week-old (1w), 3-weeks-old (3w) and adult synovium. **(B)** Uniform manifold approximation and projection (UMAP) visualisation of single-cell RNA sequencing (scRNAseq) data from total synovial cells across the developmental stages analysed. **(C)** UMAP visualisation of total synovial cells stratified by developmental stage. **(D)** Heatmap and **(E)** bubble plot showing pseudo-bulk expression of key marker genes defining all synovial cell clusters. **(F)** Quantification of synovial cell cluster distribution across developmental stages. **(G)** Heatmap showing pseudo-bulk expression of top marker genes defining macrophage and monocyte clusters. **(H)** UMAP visualisation of macrophages and monocytes in adult males and females. **(I)** Differentially expressed genes (DEGs) in VSIG4⁺ lining macrophages comparing adult versus 1 week-old and adult versus 3 weeks-old mice. **(J)** DEG analysis of adult VSIG4⁺ lining macrophages between both sexes. Data are presented as mean ± SD. Statistical significance was determined using the Kruskal– Wallis test with Dunn’s correction for multiple comparisons, relative to 1w. *p<0.05, **p<0.01.

**Figure S3.**
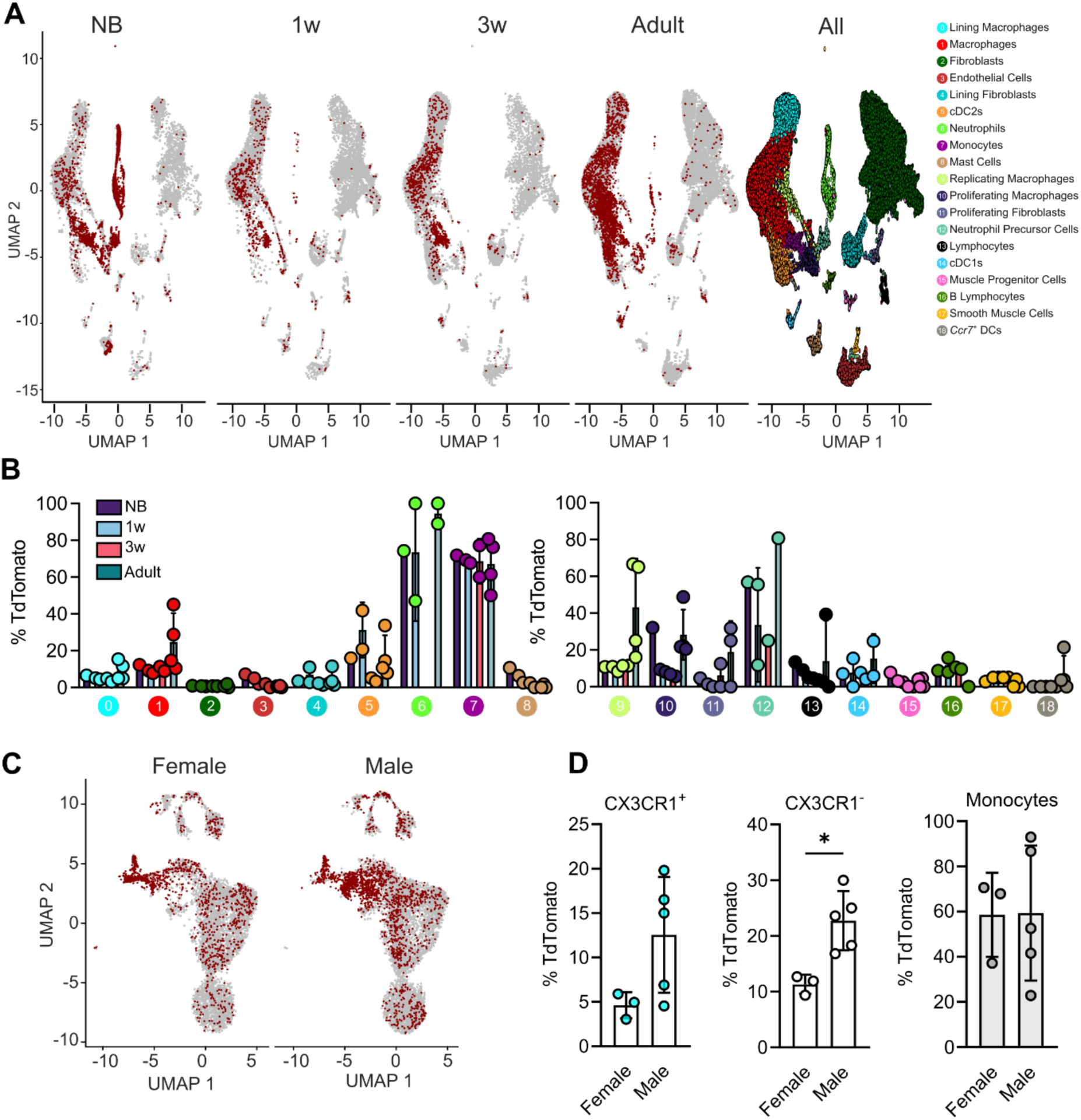
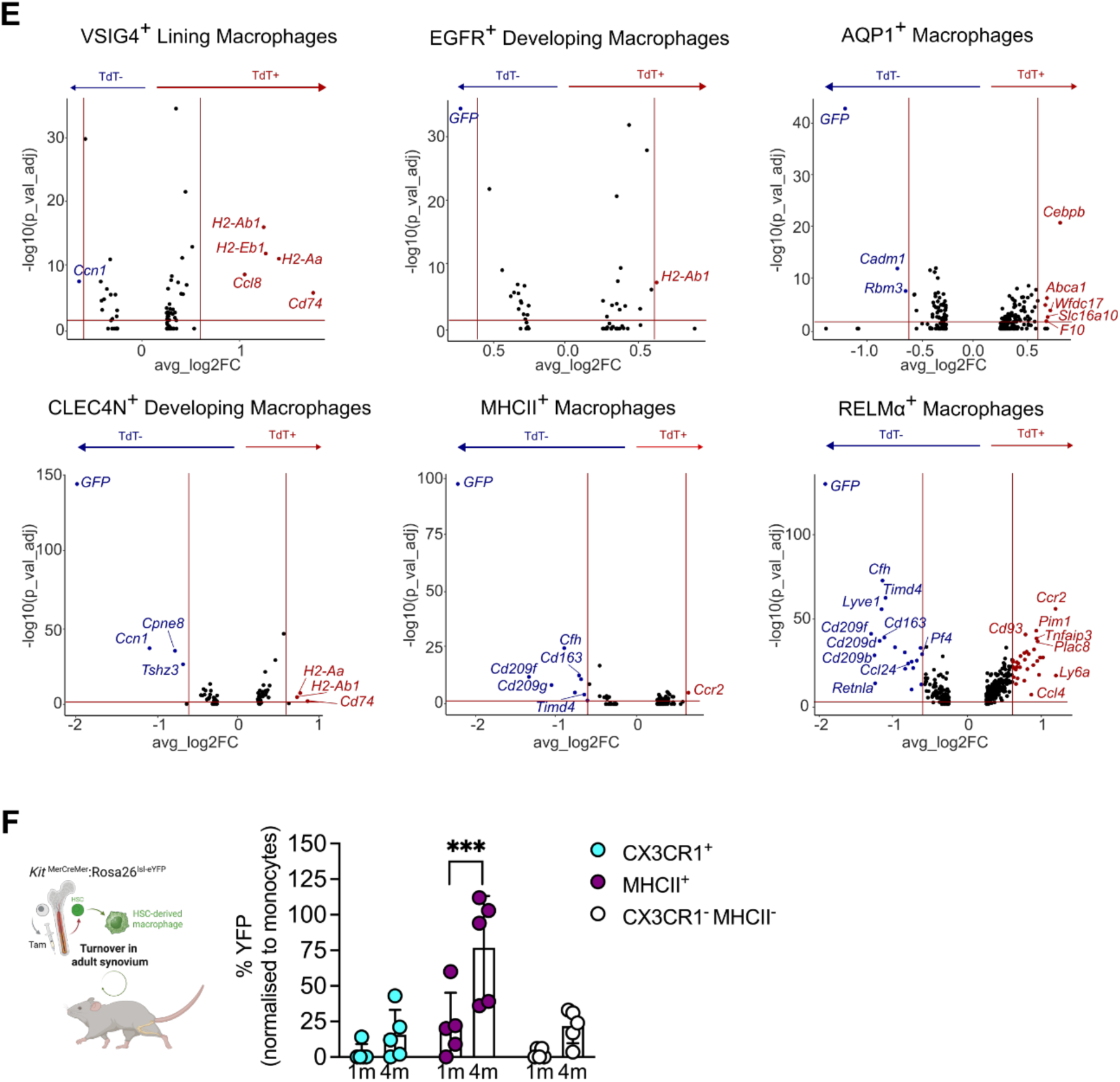
Synovial lining macrophages receive limited input from monocytes. **(A)** Uniform manifold approximation and projection (UMAP) overlay of single-cell RNA sequencing data (scRNAseq) showing *tdTomato* expression for all synovial cell types. **(B)** Abundance of tdTomato⁺ cells in synovial populations. **(C)** UMAP visualisation comparing *tdTomato* expression between male and female adult samples. **(D)** Flow cytometry analysis of tdTomato⁺ CX3CR1⁺ lining and CX3CR1⁻ sublining macrophages in adult male and female mice. Ly6C^high^ blood monocytes served as a positive control. **(E)** Differential gene expression analysis between *tdTomato*⁺ and *tdTomato*⁻ cells for different macrophage subsets. **(F)** Synovial macrophage turnover from bone marrow HSCs. (Left) Schematic of the *Kit*^MerCreMer^:Rosa26^lsl-eYFP^ fate mapping model. In adult mice, tamoxifen treatment labels *Kit*-expressing cells, including bone marrow HSCs. (Right) Flow cytometric quantification of YFP labelling in CX3CR1⁺ lining and CX3CR1⁻ MHCII⁻ and CX3CR1^-^ MHCII⁺ sublining macrophages 1 and 4 months after injection. For scRNAseq, each dot represents a pool of mice representing a mixed biological replicate as described in the Material and Methods section. For flow cytometry, each dot represents an individual mouse. Data are presented as mean ± SD. Statistical significance was determined using the Mann–Whitney test for pairwise comparisons. *p<0.05, ***p<0.001.

**Figure S4.**
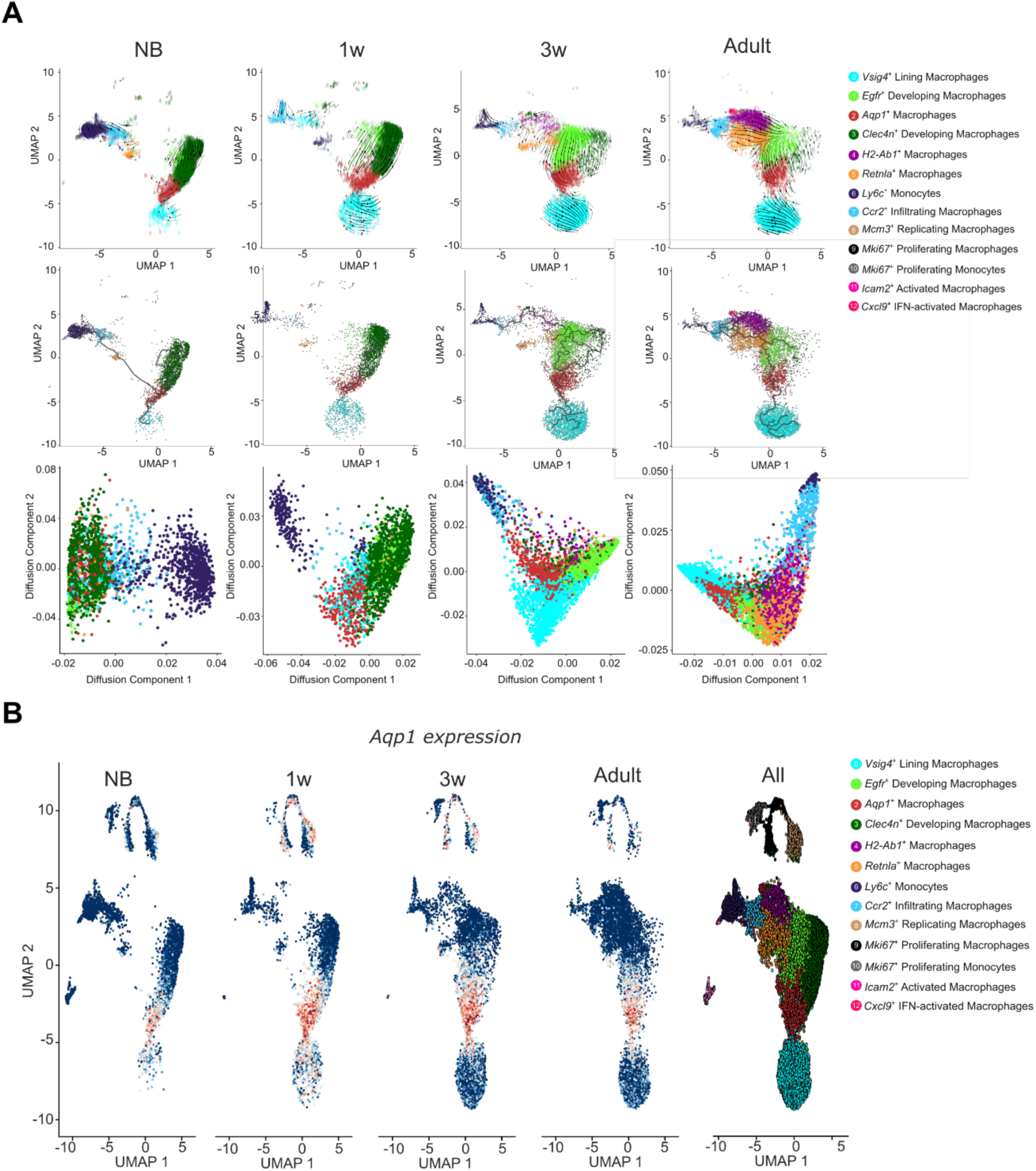
Inference of developmental relationship between *Aqp1*^+^ and lining macrophages. **(A)** Trajectory analyses to infer developmental relationships between synovial macrophages during postnatal development. Alternative algorithms were used to complement RNA velocity trajectories (Figure 4B). (Top) RNA velocity analysis visualised on uniform manifold approximation and projection (UMAP), where arrow direction indicates inferred cell trajectory based on spliced versus unspliced RNA counts. (Middle) Monocle trajectory projection overlaid on UMAP. (Bottom) Diffusion maps displaying predicted transitions between macrophage states. **(B)** UMAP visualisation of macrophages at the indicate stages, with *Aqp1* expression overlaid.

**Figure S5.**
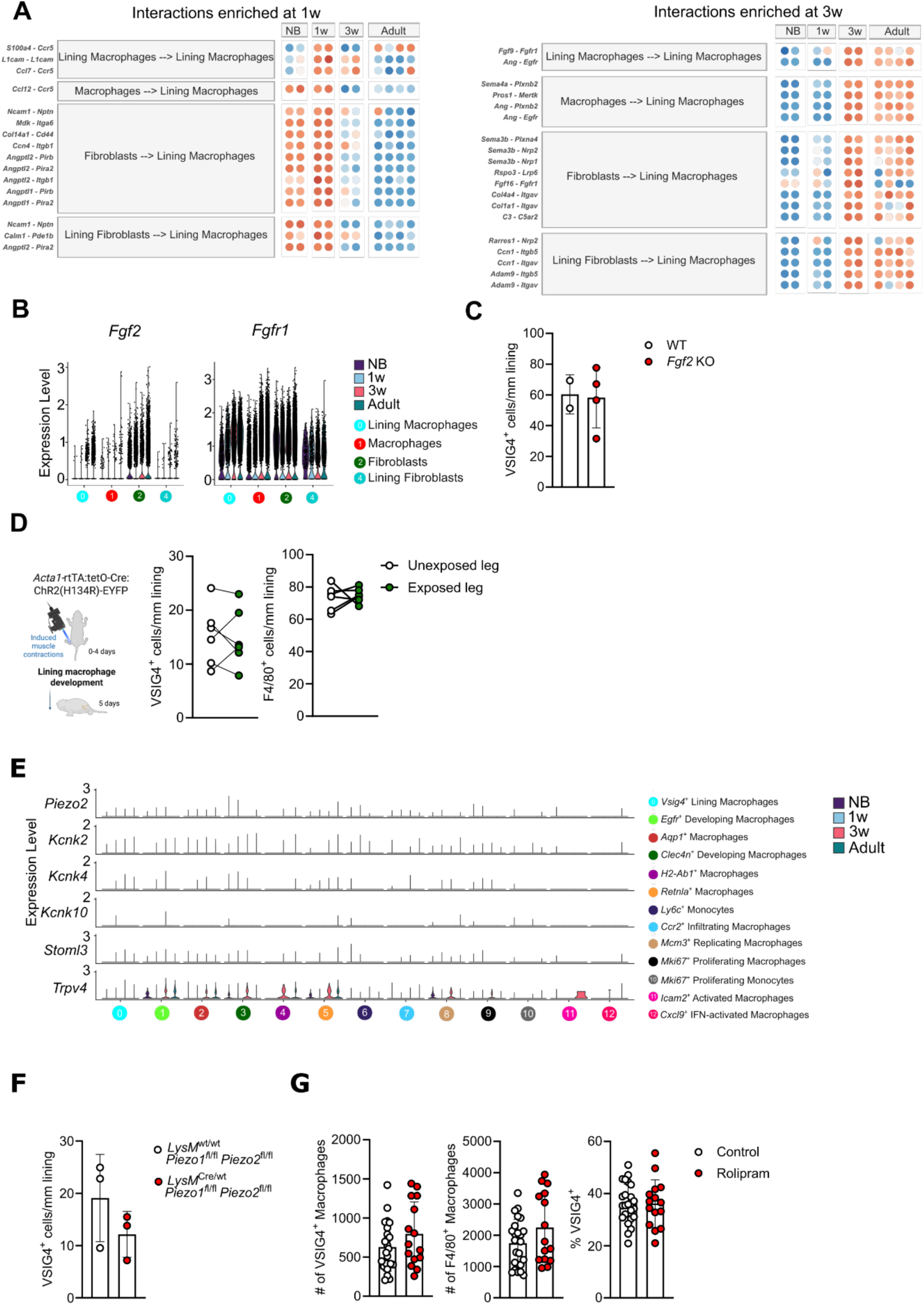
Involvement of mechanical loading and mechanosensing in lining macrophage development. **(A)** Identification of candidate signals predicted to instruct lining macrophage development based on MultiNicheNet analysis of scRNAseq data. Bubble plots show the top enriched pathways at 1 week (Left) and 3 weeks (Right). **(B, C)** FGF2 signalling is not involved in lining macrophage development. **(B)** Expression of *Fgf2* and *Fgfr1* by synovial macrophages and fibroblasts at the distinct developmental stages. **(C)** Lining macrophage development in *Fgf2* knockout mice. The density of VSIG4^+^ lining macrophages was determined by confocal imaging for 3-weeks-old mice. **(D)** Impact of optogenetically induced neonatal leg muscle contractions on lining macrophage development. (Left) Schematic of the experimental approach. *Acta1-rtTA*^tetO-Cre^:*ChR2-V5*^fl^ mice were used, in which skeletal muscles express a light-sensitive channelrhodopsin. Muscle contractions were induced daily in pups for the first four days of life. The contralateral limb was not exposed to light and served as a control. (Right) The effects of muscle contractions on lining macrophage density were determined by confocal imaging at 5 days of age (right). **(E)** Mechanoreceptor gene expression by synovial macrophages during postnatal development. **(F)** Synovial macrophage development in mice with loss of function in PIEZO1 and PIEZO2 signalling. Myeloid-specific *Piezo1* and *Piezo2* knockout mice (*LysM*^Cre^:*Piezo1*^fl/fl^ *Piezo2*^fl/fl^) were analysed at 3 weeks of age. Confocal imaging was used to quantify the density of VSIG4⁺ macrophages in the synovial lining. **(G)** Involvement of the Phosphodiesterase-4 (PDE4) pathway in synovial macrophage development. PDE4 activity was inhibited by intra-peritoneal administration of Rolipram (0.5 μg per gram of body weight) at 3, 5 and 7 days of age. The abundance of total and VSIG4^+^ macrophages as well as the frequency of VSIG4^+^ cells within synovial macrophages were determined in 3-weeks-old mice by flow cytometry. For imaging analyses, each dot represents an individual mouse, with quantification based on 2 to 3 sections per animal. For flow cytometry, each dot represents an individual mouse. Statistical significance was determined using the Mann–Whitney test (unpaired) or Wilcoxon test (paired). *p<0.05.

**Supplementary Table 1:**
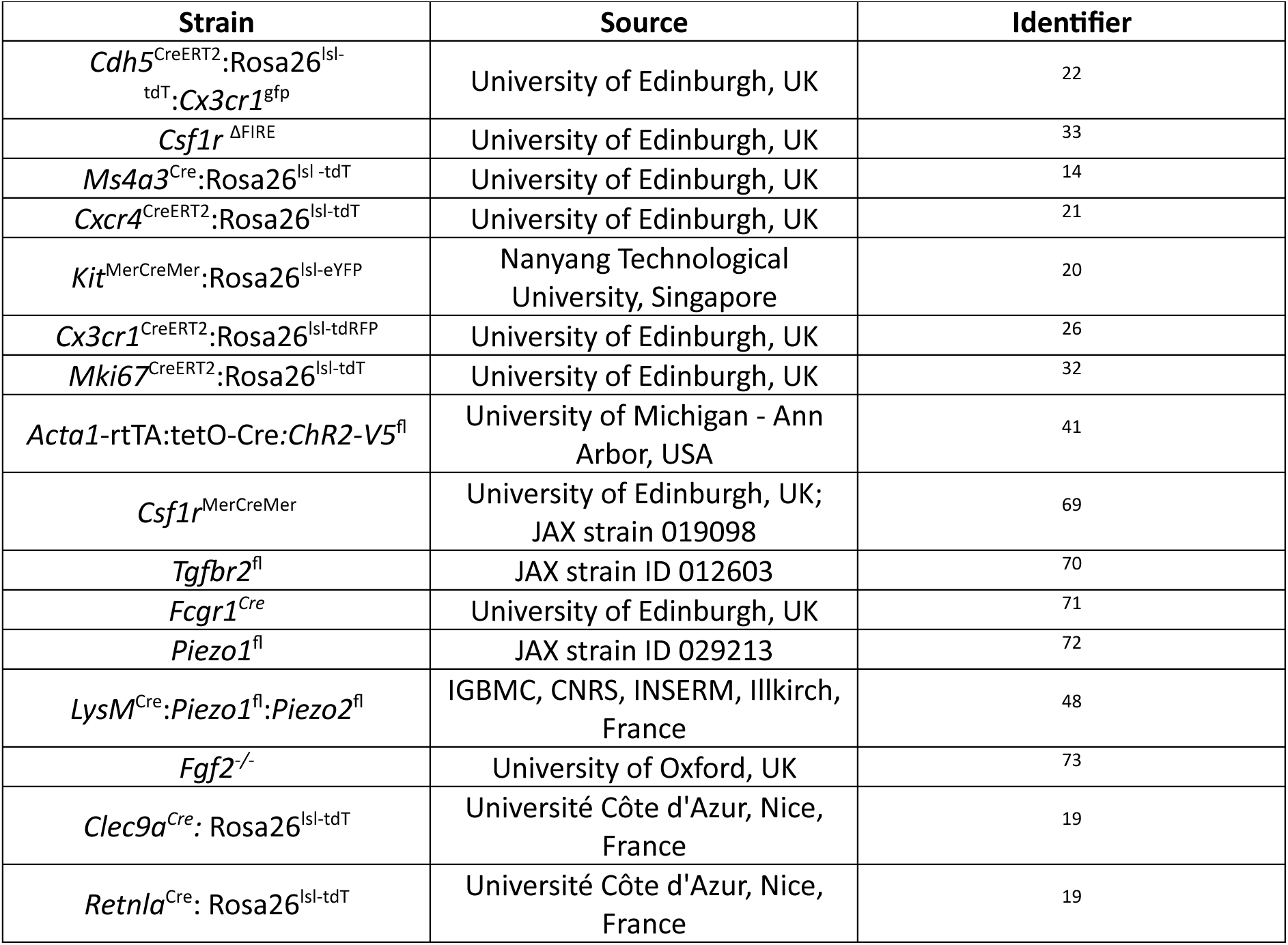
Mouse strains.

**Supplementary Table 2:**
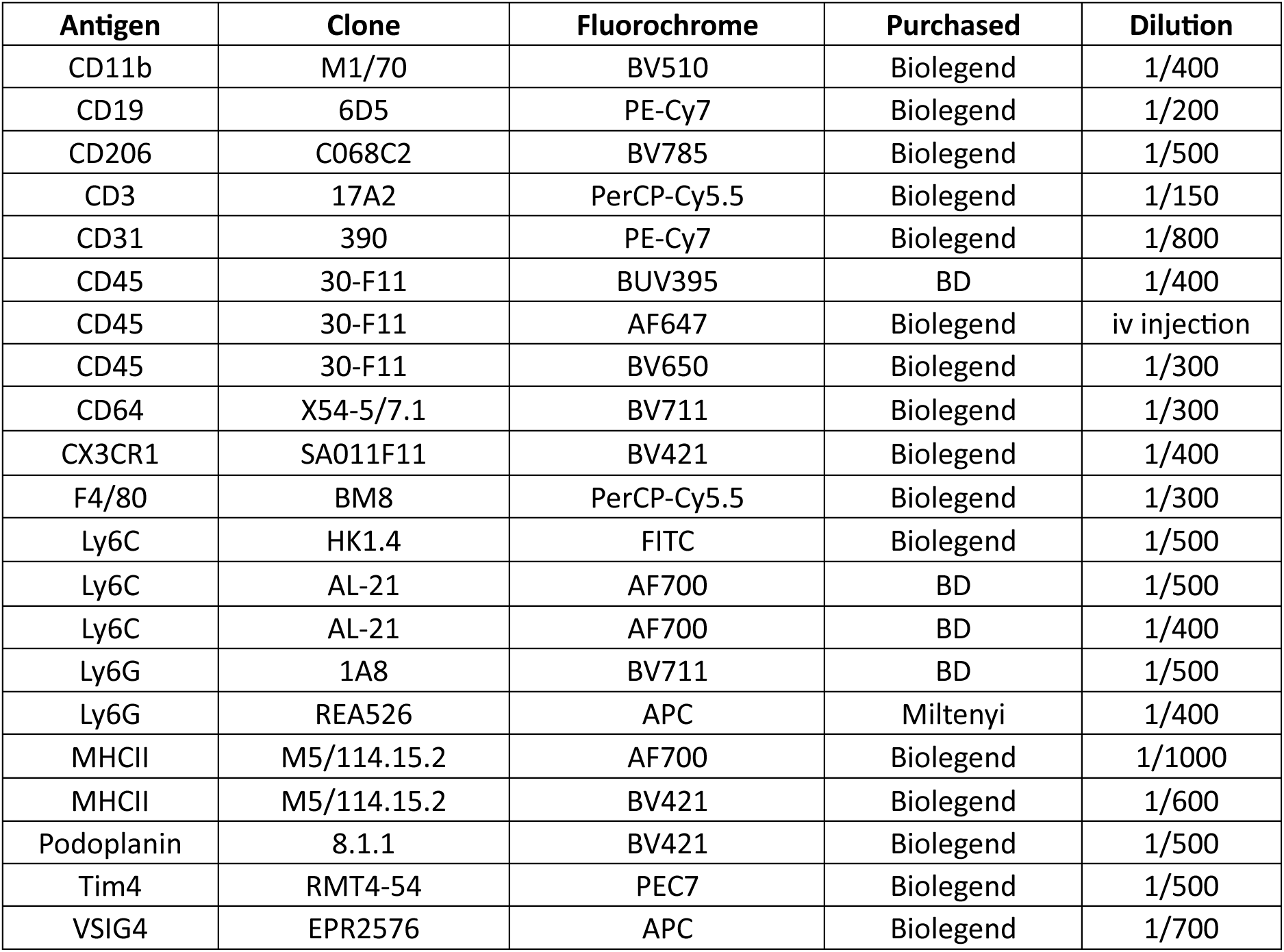
Antibodies for flow cytometry.

**Supplementary Table 3:** Loading strategy for scRNAseq.

| Condition | Number of mice (Sex) | Number of litters | Oligo labelling by | Immune cells loaded | Fibroblasts/Endothelial cells loaded |
| --- | --- | --- | --- | --- | --- |
| Adult 1 | 8 (4M, 4F) | 2 | Sex | 40 000 | 20 000 |
| Adult 2 | 11 (9M, 2F) | 2 | Litter | 30 000 | 35 000 |
| Newborn | 19 (Mix) | 3 | n/a | 1000 | 10 000 |
| 3 weeks | 20 (Mix) | 4 | Sex | 40 000 | 20 000 |
| 1 week | 20 (Mix) | 3 | Sex | 33 000 | 27 000 |
| BBP | 9 (7M, 2F) | 2 | Litter | 30 000 | 35 000 |

**Supplementary Table 4:**
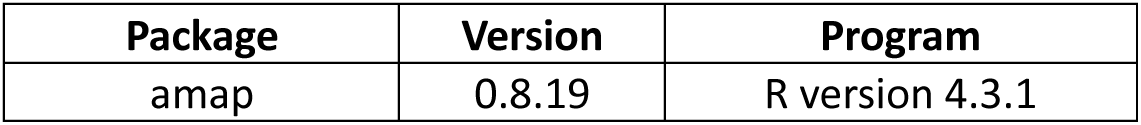
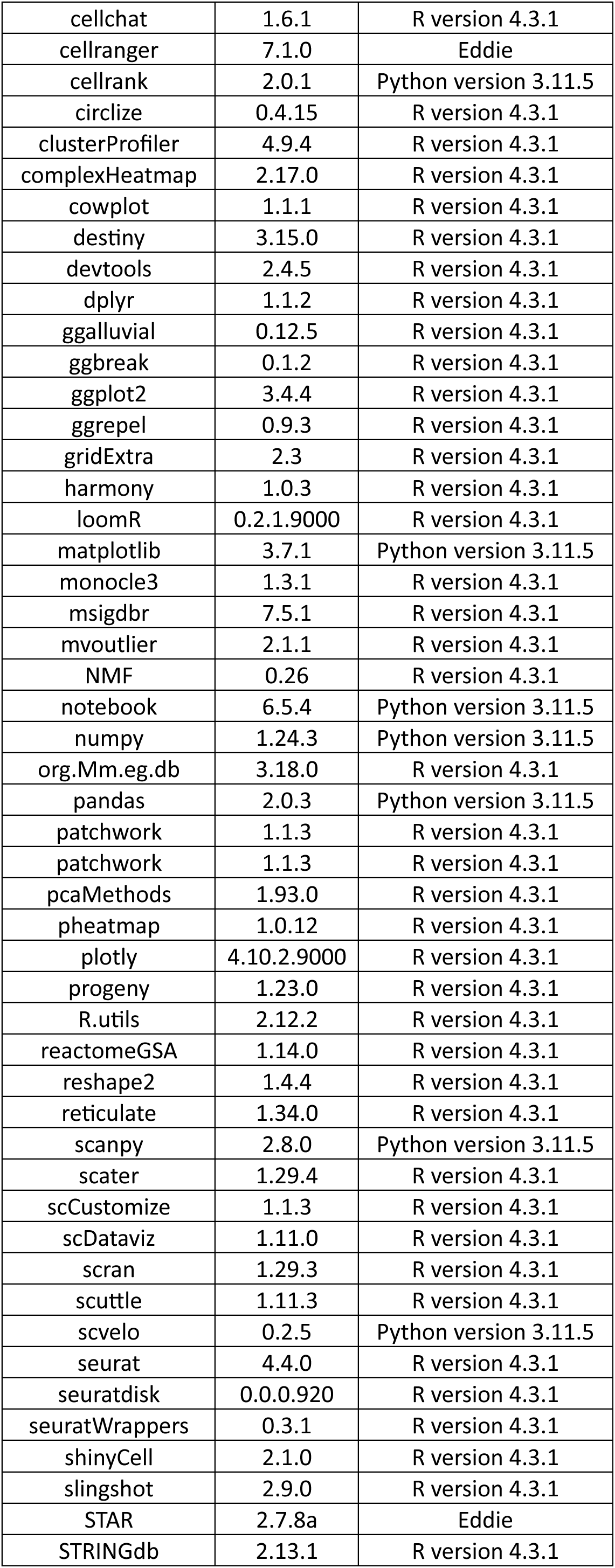
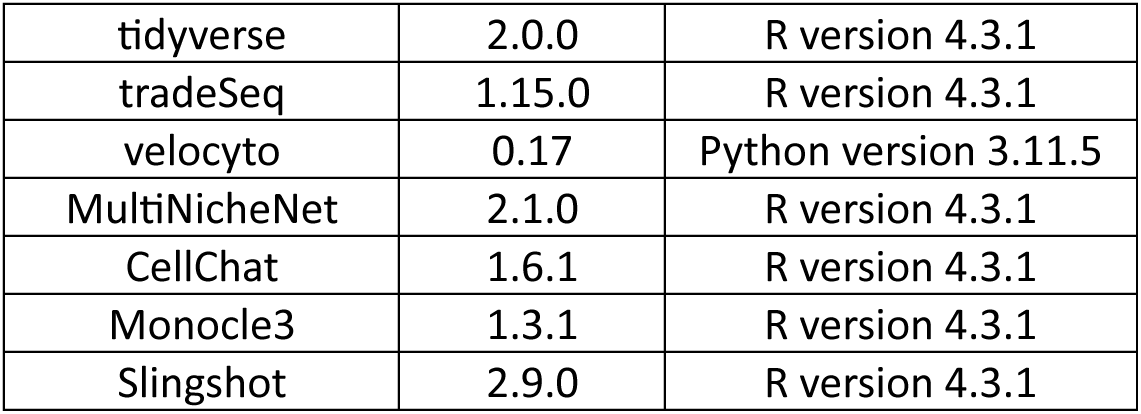
Software packages for scRNAseq analysis.

**Supplementary Table 5:** Primary antibodies for immunofluorescence.

| Antigen | Species raised | Clone | Fluorophore | Purchased | Dilution |
| --- | --- | --- | --- | --- | --- |
| F4/80 | Rat | Cl:A3-1 | APC | Abcam | 1/400 |
| VSIG4 | Rabbit | EPR22576-79 | n/a | Abcam | 1/500 |
| CD140a (PDGFRα) | Rat | APA5 | n/a | Invitrogen | 1/100 |
| IBA1 | Goat | Poly | n/a | Invitrogen | 1/500 |
| AQP1 | Rabbit | EPR11588(B) | n/a | Abcam | 1/200 |

**Supplementary Table 6:**
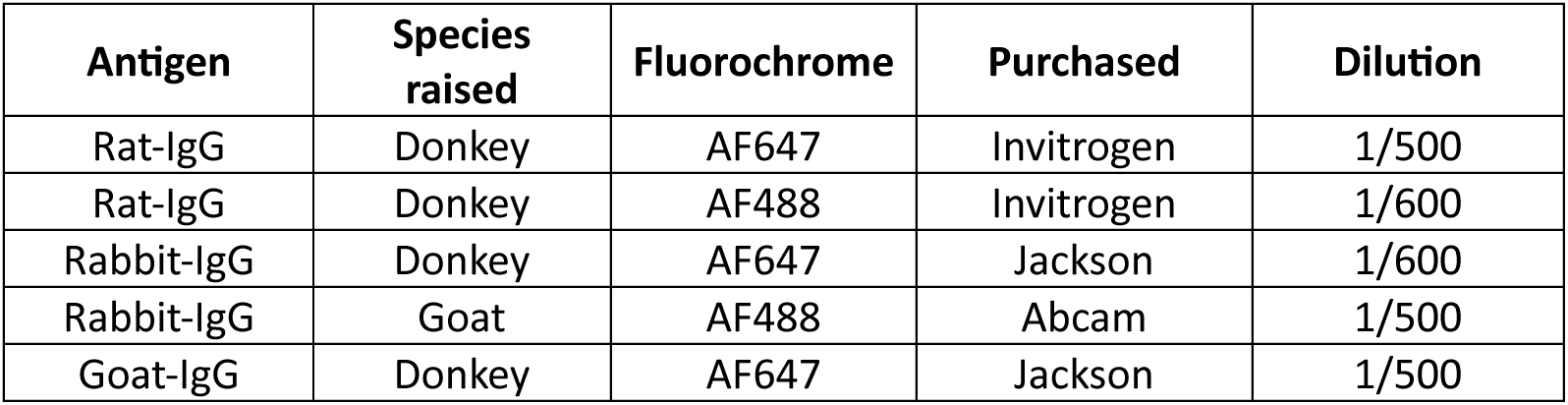
Secondary antibodies for immunofluorescence.

